# Rewired m6A methylation of promoter antisense RNAs in Alzheimer’s disease regulates global gene transcription in the 3D nucleome

**DOI:** 10.1101/2025.03.22.644756

**Authors:** Benxia Hu, Yuqiang Shi, Feng Xiong, Yi-Ting Chen, Xiaoyu Zhu, Elisa Carrillo, Xingzhao Wen, Nathan Drolet, Chetan Rajpurohit, Xiangmin Xu, Dung-Fang Lee, Claudio Soto, Sheng Zhong, Vasanthi Jayaraman, Hui Zheng, Wenbo Li

## Abstract

N^6^-methyladenosine (m6A) is the most prevalent internal RNA modification that can impact mRNA expression post-transcriptionally. Recent progress indicates that m6A also acts on nuclear or chromatin-associated RNAs to impact transcriptional and epigenetic processes. However, the landscapes and functional roles of m6A in human brains and neurodegenerative diseases, including Alzheimer’s disease (AD), have been under-explored. Here, we examined RNA m6A methylome using total RNA-seq and meRIP-seq in middle frontal cortex tissues of post-mortem human brains from individuals with AD and age-matched counterparts. Our results revealed AD-associated alteration of m6A methylation on both mRNAs and various noncoding RNAs. Notably, a series of <u>p</u>romoter <u>a</u>ntisense RNAs (paRNAs) displayed cell-type-specific expression and changes in AD, including one produced adjacent to the *MAPT* locus that encodes the Tau protein. We found that *MAPT-paRNA* is enriched in neurons, and m6A positively controls its expression. In iPSC-derived human excitatory neurons, *MAPT-paRNA* promotes expression of hundreds of genes related to neuronal and synaptic functions, including a key AD resilience gene *MEF2C*, and plays a neuroprotective role against excitotoxicity. By examining RNA-DNA interactome in the three-dimensional (3D) nuclei of human brains, we demonstrated that brain paRNAs can interact with both *cis*- and *trans*-chromosomal target genes to impact their transcription. These data together reveal previously unexplored landscapes and functions of noncoding RNAs and m6A methylome in brain gene regulation, neuronal survival and AD pathogenesis.

## Introduction

Alzheimer’s disease (AD) is the most common cause of dementia (60–70% of all cases), and affects 5% of the population over 60 years (or approximately 1% of the total population)^1^. AD is pathologically characterized by massive neuronal loss, the presence of amyloid-beta (Aβ) plaques, and Tau neurofibrillary tangles (NFTs) in the brain^2,3^. Despite extensive studies of AD, there are still limited therapeutic options that can effectively treat this devastating disease. Identifying new molecular players in AD pathogenesis is important to offer further insights and potential targets for combating this disease.

In the past decade, studies of RNA chemical modifications have founded a field that is referred to as “epitranscriptomics”^4,5^. Among these RNA modifications, N^6^-methyladenosine (m6A) is the most abundant internal RNA methylation in mammalian cells, and it can impact mRNA stability, trafficking, and translation^4,5^. Analogous to the mechanism of epigenetics, biological functions of m6A are considered to be mediated by specific proteins that write, read, and erase this mark, which are referred to as m6A writers (e.g., METTL3), readers (e.g., YTH proteins), and erasers (e.g., FTO and ALKBH5), respectively^4,5^. In addition to post-transcriptional roles on mRNAs, m6A methylation is increasingly realized to bear transcriptional roles by modifying nuclear or chromatin-associated RNAs to control gene transcription via functionally cross-talking to epigenetic processes^6–10^. Examples include m6A regulation of *Xist* lncRNA^8,11^, retrotransposon RNAs^10,12–14^, or enhancer RNAs^7,15^. These results have provided a new perspective to study the roles of m6A in gene regulation in development and diseases, which encompasses both transcriptional and post-transcriptional processes. Interestingly, m6A methylation seems to play unique functions in the brain. For example, in both mice and human brains, m6A landscapes display a pattern distinctive from other tissues^16^. In murine models, conditional depletion of genes encoding a component of the m6A methyltransferase complex, Mettl14, or a cytoplasmic m6A reader protein, Ythdf1, severely affected neurogenesis and cognitive functions^17,18^. However, despite efforts^19^, the landscapes and functional roles of m6A or other epitranscriptomic modifications in human brains and AD remain minimally explored.

Regulatory long noncoding RNAs (lncRNAs) are pervasively produced in the human genome^20,21^. A large fraction of these transcripts originate from divergent transcription at promoters of active protein-coding genes (>60% in human ESCs^22^), which are interchangeably referred to as promoter antisense RNAs (paRNAs) or PTOMPTs^23–27^. Some studies reported that paRNAs can regulate the expression of mRNAs divergently produced from the same promoters^27,28^. Actually, deregulation of lncRNAs has been long observed in AD^29–31^. An early example is an antisense RNA produced within the *BACE1* gene locus that was proposed to regulate *BACE1* expression to impact AD disease progression^32^. High resolution transcriptomes in AD brains have now permitted rapid discovery of enormous ncRNAs^30,31^. A recent work suggested that AD-associated <u>n</u>atural <u>a</u>ntisense transcripts (NAT), some of which are paRNAs, can play roles in the translational control of crucial disease proteins including MAPT/Tau to impact AD pathogenesis^33^. However, another study reported inconsistent results^34^. These studies together indicated that the roles of lncRNAs are still under-studied in human AD brains, and their mechanisms of action are poorly understood.

Here, we characterized RNA m6A methylome from individuals diagnosed with AD and from cognitively normal age-matched individuals. We uncovered a large number of m6A-modified RNAs in the human brain, including both mRNAs and ncRNAs, many of which were previously unannotated. RNA m6A methylome in AD brains displays alteration as compared to controls, and there is positive correlation between its signals and expression levels of mRNA and ncRNAs. Subsequently, we focused on paRNAs and used induced pluripotent stem cell (iPSC)- derived human excitatory neurons (i3Neurons) to functionally investigate regulatory roles of an m6A-modified *MAPT-paRNA* transcribed next to *MAPT*, a key AD-associated disease locus. Our data revealed that *MAPT-paRNA* shows neuron-preferred expression and is upregulated in AD. It promotes the expression of many neuronal and synaptic genes via navigating the three-dimensional (3D) genome organization, which is required for neuronal survival under excitotoxic conditions. This work unraveled previously unexplored landscapes and the role of chemical modification on AD-associated ncRNAs, linking epitranscriptome to gene transcriptional control in the 3D genome and AD pathogenesis.

## Results

### The transcriptome and m6A epitranscriptome of human AD brains on coding and noncoding RNAs

From a cohort of individuals diagnosed with sporadic AD (n = 6; mean age = 66) and from cognitively normal age-matched counterparts (n = 6; mean age = 65, hereafter referred to as Normal), we collected the middle frontal cortex tissue (mFC), a region pathologically affected in AD^35^. Subsequent to autopsy, neuropathologic examination revealed high levels of phosphorylated Tau neurofibrillary tangles and amyloid-β burden (**Supplementary Table 1**). We analyzed the transcriptome by strand-specific ribo-depleted total RNA-seq, with the aim of examining both mRNAs and various ncRNAs. RNA m6A methylome was examined by m6A antibody-based immunoprecipitation (MeRIP-seq) (**see methods**). A summary of our sequencing datasets can be found in **Supplementary Table 2**.

Analysis of total RNA-seq data identified 416 and 519 genes significantly upregulated and downregulated in AD as compared to Normal groups, respectively (**Supplementary Fig. 1A**, gene lists in **Supplementary Table 3**). These changes are consistent with a previous study that conducted total RNA-seq in the lateral temporal lobe (LTL) region of AD/Normal human brains^36^ (**Supplementary Fig. 1B,C**). Gene Ontology (GO) analysis of differentially expressed genes (DEGs) showed that AD-upregulated genes are related to development, while the AD-downregulated genes are enriched in neuronal/synaptic functions (**Supplementary Fig. 1D,E,** and **Supplementary Table 4**). This is in accord with the knowledge that neuronal/synaptic genes are reduced in AD^37^. This also suggests that tissue-level gene deregulation that we found here largely reflects changes in neurons^38^. Several representative examples of deregulated genes are shown in **Supplementary Fig. 1F**. *YAP1*, a gene encoding a key transcription regulator, was induced in AD, whereas AD-downregulated genes are exemplified by neuronal transcriptional factor gene *NEUROD6* and those encoding glutamate decarboxylases essential for inhibitory neurotransmitter gamma-aminobutyric acid (GABA) synthesis, e.g., *GAD1*/*GAD2* (**Supplementary Fig. 1F**).

To examine the changes of RNA m6A methylome in AD versus Normal brains, we called strand-specific m6A peaks on total RNA-Seq/MeRIP-seq datasets using MACS3^39,40^, and identified approximately 52,661 and 55,754 m6A peaks in Normal and AD mFC tissues, respectively (**Supplementary Table 5**). A majority of peaks locate in the non-protein-coding portions of the genome, i.e., around 60% were in intergenic regions, introns or promoter upstream regions (−2kb to transcription start sites) (**Fig. 1A**). The remaining peaks overlapped with protein-coding mRNA sequences (CDSs), and 5’ or 3′ untranslated regions (UTRs) (**Fig. 1A**). This is consistent with the higher sensitivity of ribo-depletion strategy in RNA-seq to detect various ncRNAs. We compared the features of m6A peaks in Normal and AD conditions, and found them to be overall similar: they enrich the canonical RRACH motif, bearing comparable peak length (Normal: 236bp v.s. AD: 227bp) and average GC contents (0.47 v.s. 0.45) (**Supplementary Fig. 2A,B,C**). The genomic distribution of m6A signals is quite similar between RNAs from Normal and AD brains, too (**Supplementary Fig. 2D**). These results indicate that AD elicits a quantitative rewiring of the RNA m6A landscape in the human brain, at least in the mFC region.

**Figure 1.**
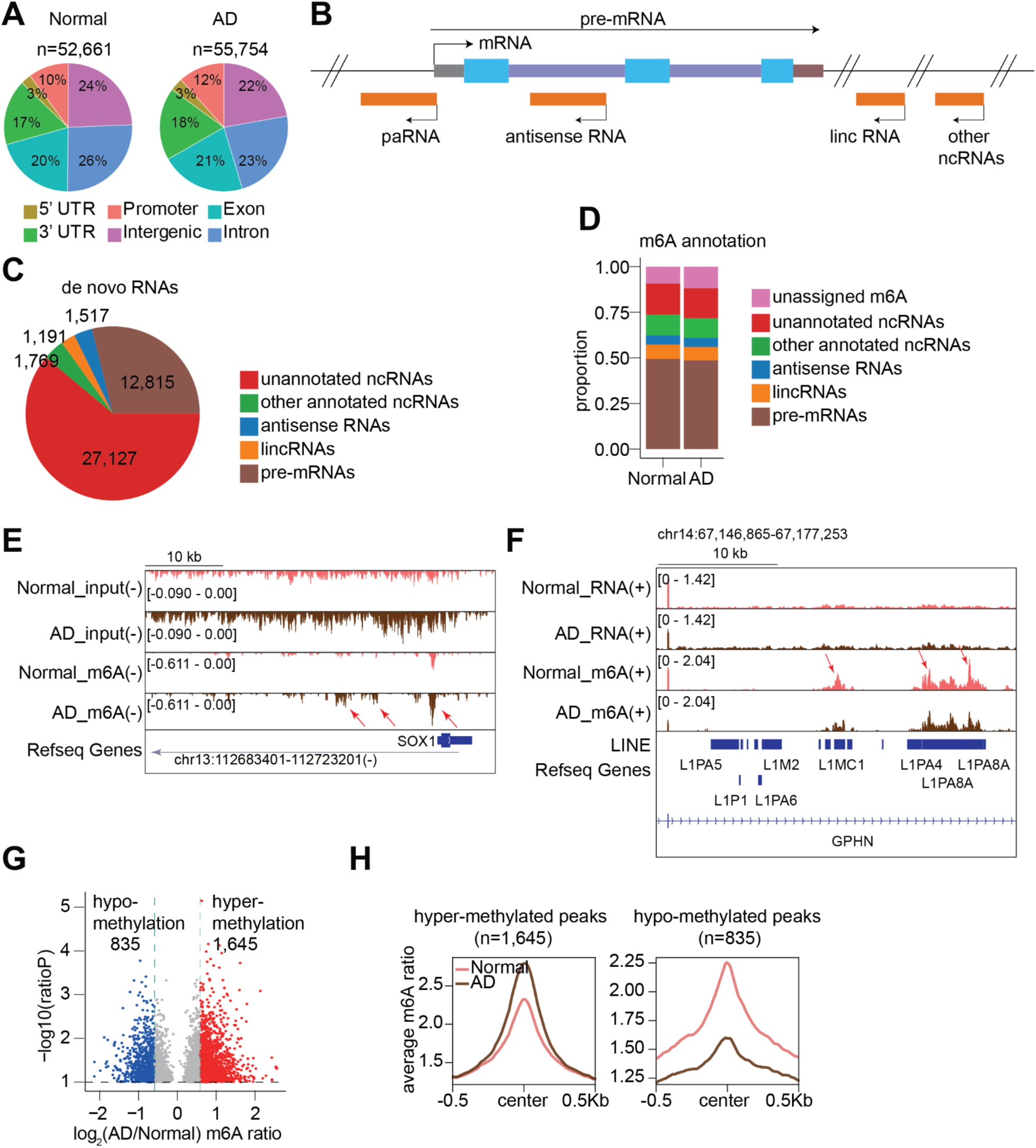
m6A methylome on coding and noncoding RNAs in the mFC regions of human brains. **A**. Piecharts showing the m6A peak distribution based on genomic locations in post-mortem human mFC tissues from Normal and AD donors, respectively. **B.** A model illustrating major categories of *de novo* identified transcripts, including pre-mRNAs and ncRNAs. **C**. A piechart showing the numbers of transcripts identified by the de novo calling. **D**. A barplot showing the assignment of m6A peaks to various *de novo* called transcripts. **E.** Genome browser tracks showing an example of ncRNA (a promoter antisense RNA from the *SOX1* locus) having m6A peaks in human Normal and AD brains. Input and m6A represent the average signals of RNA-seq and meRIP-seq datasets in this work, respectively. The arrows indicate m6A peaks. **F.** Similar to E, example tracks showing intronic L1 elements overlap m6A peaks in the intron of *GPHN* gene. (+) and (-) indicate the Watson and Crick strands, respectively. **G.** A volcano plot showing differential m6A peaks between Normal and AD brains. Red and blue dots represent hyper- and hypo-methylated m6A sites in AD. **H**. Metaplots showing the aggregated m6A ratios of hyper- and hypo-methylated m6A sites.

A large number of m6A peaks locating in the noncoding regions is consistent with recent work about m6A on regulatory ncRNAs^7,8,12,15,41^. Annotation of m6A peaks using genomic locations is not sufficient to precisely associate them to specific noncoding RNA transcripts (**Fig. 1A,B**). We therefore need to first define a complete catalog of transcripts and their genomic coordinates in the human brain by a *de novo* transcript calling method based on Hidden Markov Model (HMM) ^42,43^. Of the *de novo* identified transcripts, 12,815 overlap protein-coding genes and are thus defined as pre-mRNAs in this work (**Fig. 1B,C**). There are 31,604 ncRNA transcripts in the human brain (**Fig. 1C**), including 27,127 previously unannotated RNAs in the human transcriptome. Of the remaining, 1,191 overlap annotated lncRNAs, 1,517 overlap annotated antisense RNAs, and 1,769 overlap other annotated RNAs (snoRNAs, snRNAs, pseudo genes) in gencode v19 (**Fig. 1C**). Approximately 90% of all m6A peaks can be assigned to these various *de novo* identified RNAs (**Supplementary Fig. 2E,** and **Fig. 1D**). The ∼10% unassigned peaks are largely due to the fact that some RNAs are not identified as independent transcripts by the *de novo* calling algorithm and thus m6A peaks on them are not assigned. As an example, of the 27,127 previously unannotated transcripts (**Fig. 1C,D**), an antisense RNA generated from the *SOX1* gene promoter shows prominent m6A peaks (**Fig. 1E**). About 50% of m6A peaks were assigned to the sense strand of pre-mRNAs, including exons, introns, 5’UTR and 3’UTR (**Fig. 1D,F**). Among these, interestingly, over ten-thousand peaks appear in introns and often overlap transposable elements, in particular, intronic L1 elements (**Fig. 1F**, and **Supplementary Fig. 2F,G,H**). This is reminiscent of our previous work showing that many intronic L1 RNAs bear high m6A methylation and play roles in gene transcriptional control^12^. Collectively, at the transcriptome level, 60-70% of all m6A peaks are located in ncRNAs or introns as opposed to mRNA regions (UTRs and exons), and the distribution pattern is largely similar in AD versus Normal (**Supplementary Fig. 2H**). These results provided a comprehensive m6A RNA methylome on mRNAs and ncRNAs in human Normal and AD brains, serving as a foundation to study the functions of these m6A signals in gene expression regulation and AD biology.

### The alteration of m6A epitranscriptome in the human AD brains

By calculating differential m6A methylation (**see methods**), we identified 835 and 1,645 <u>d</u>ifferential <u>m</u>6A <u>p</u>eaks (DMPs) showing hypo- and hyper-m6A-methylation in AD, respectively (**Fig. 1G**). Here we defined m6A methylation levels as m6A ratios, the signal division between meRIP-seq and RNA-seq in each peak. The changes in m6A ratios at the differential peaks can be seen by meta-analysis (**Fig. 1H**), and the motifs in these regions remain canonical RRACH (**Supplementary Fig. 3A**). A breakdown shows that DMPs overall occurred proportionally to the total peak distribution, with about 40% of DMPs on pre-mRNAs (including exons, UTRs and introns) and the other ∼60% of DMPs are observed to be from ncRNAs (**Fig. 1D**, and **Supplementary Fig. 3B**).

Previous m6A studies have shown correlation between changes of m6A and mRNA expression levels^4,44^. We examined their correlation in AD brains by examining mRNAs bearing DMPs on their exons or UTRs. By cumulative fraction analysis, we observed a positive correlation between the two, i.e., m6A hyper-methylation is associated with mRNA expression increase in AD, whereas hypo-methylation is associated with decrease (**Supplementary Fig. 3C,D**). Interestingly, GO enrichment analyses of genes whose mRNAs display hyper-methylation show terms related to transcriptional activities, whereas gene mRNAs with hypo-methylated peaks are more relevant to synapse regulation (**Supplementary Fig. 3E,F**). Because of the known expression reduction of synapse related genes in AD^37^, this result suggests that hypo-methylation of their mRNAs is associated with expression reduction during AD pathogenesis. We showed two representative examples of m6A peaks in the 3’UTR regions that were hyper- or hypo-methylated in AD compared to Normal, which correlates with their respective mRNA expression changes (**Supplementary Fig. 3G,H**). We conducted similar analysis for ncRNAs, and observed that m6A changes are also positively correlated with ncRNA expression alteration in AD, i.e., hyper-methylation is associated with ncRNA expression increase, whereas hypo-methylation with decrease (**Supplementary Fig. 4A,B**). We show two representative examples of ncRNA regions that were increased or decreased in AD compared to Normal, respectively (**Supplementary Fig. 4C,D**).

### Identification and alteration of promoter antisense RNAs in AD brains

We sought to focus on previously under-studied ncRNAs and their m6A modification in AD. Visual inspection revealed a large number of paRNAs that display m6A peaks, which interestingly include many produced from important neuronal and synaptic gene loci. For example, an m6A-marked paRNA spanning ∼18kb is generated from the *GRIN2A* locus (**Fig. 2A**), which codes for an important neurotransmitter receptor Glutamate Ionotropic Receptor NMDA Type Subunit 2A. Similarly, another m6A-marked paRNA is produced antisense to the promoter of *MAPT*, one of the most significant AD-associated loci as it encodes the Tau protein (**Fig. 2B**). Importantly, functions of paRNAs have been minimally studied or understood in brain gene regulation and AD biology. *De novo* calling identified 3,038 paRNAs in mFC brain regions and about 37-39% of them display a m6A peak in either Normal or AD brains (**Supplementary Fig. 5A**). GO term analysis showed that genes neighboring m6A-marked paRNAs are associated with neuronal and synapse functions, which are deregulated processes in AD (**Fig. 2C,D**). In comparison, mRNAs with m6A do not show such association (**Supplementary Fig. 5B**), suggesting that m6A-marked paRNAs in mFCs may have specific relevance to neuronal/synapse biology. Because we observed some paRNAs with exceptional length, we utilized public nanopore long-read direct RNA-seq as an orthogonal method to validate paRNA existence and length (from dorsolateral prefrontal cortex (DLPFC) brain region by the Rush Alzheimer’s Disease Study). Defining the lengths of paRNAs is important to assign m6A peaks to these paRNA transcripts, because the peaks can sometimes be located >10kb away from the TTSs, as exemplified in **Fig. 2A,B** (and other examples see below). Despite the low coverage and its utilization of polyA capture, long-reads RNA-seq data detected about 25% of paRNAs seen by our *de novo* calling of total RNA-seq data, and the length estimation is highly accurate (**Supplementary Fig. 5C,D**). Globally, paRNA expression levels appeared comparable in Normal and AD mFCs, but those with m6A modifications are expressed higher than those without (**Supplementary Fig. 5E,F**), suggesting a positive influence of m6A methylation on paRNAs expression.

**Figure 2.**
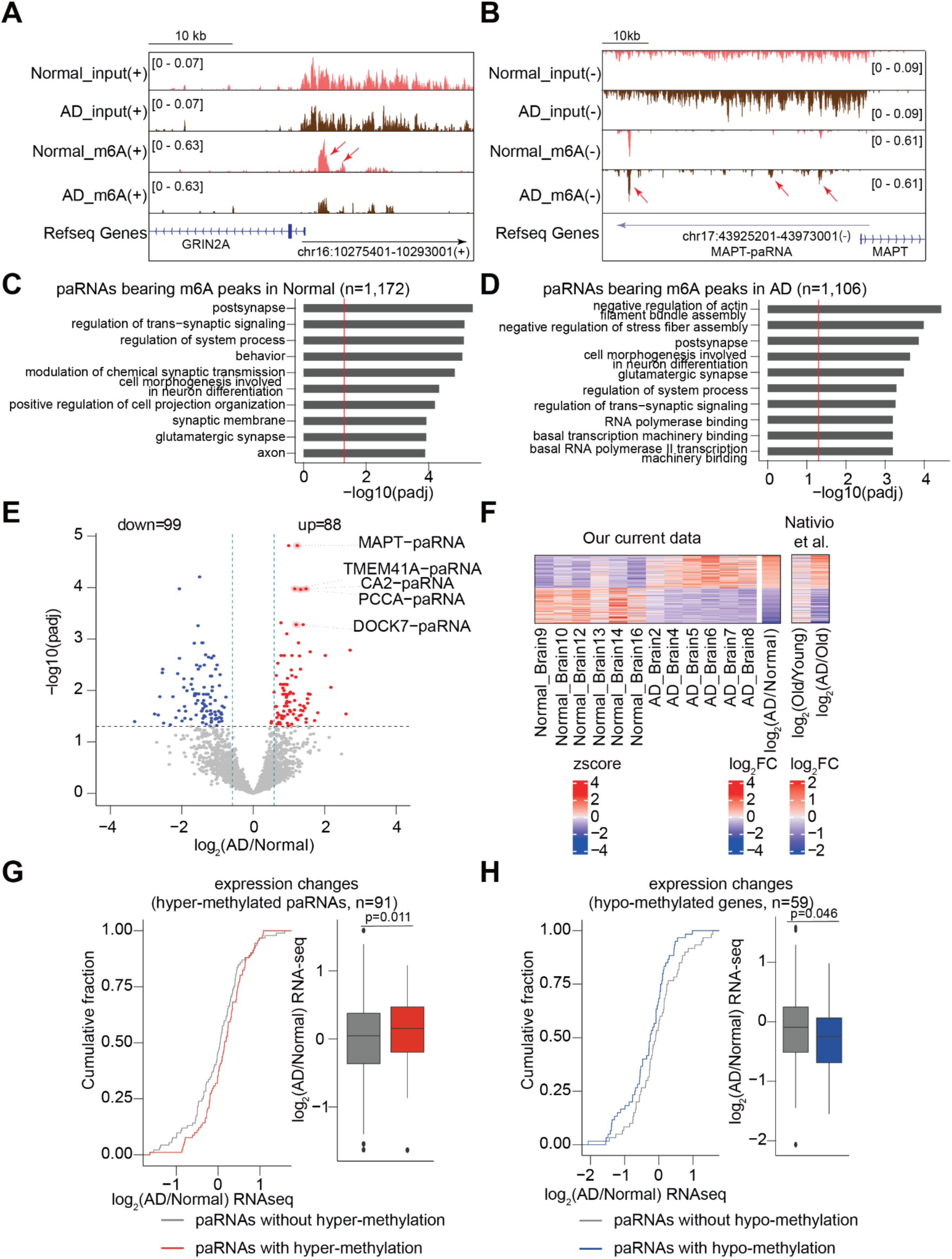
The m6A methylation and expression of paRNAs in Normal and AD human brains. **A,B**. Genome browser tracks showing the RNA-seq and m6A signals of two paRNAs (*GRIN2A-paRNA* and *MAPT-paRNA*) in the human mFC. Red arrows indicate m6A peaks. **C,D**. GO analyses for genes neighboring and sharing promoters with paRNAs. The red lines represent adjusted p values (p-adj) at 0.05. **E**. A volcano plot showing differentially expressed paRNAs between Normal and AD brains. Red and blue dots represent up-regulated and down-regulated paRNAs in AD, respectively. **F**. A heatmap showing z-score-transformed expression levels of differentially expressed paRNAs between Normal and AD brains from our samples, and from Nativio et al.^36^. **G,H.** Cumulative distribution and boxplots of paRNA expression changes with or without hyper- and hypo-m6A. P values were calculated by a two-tailed non-parametric Wilcoxon–Mann–Whitney test. Boxplots indicate the interquartile range with the central line representing the median, and the vertical lines extending to the extreme values in the group.

Differential analysis of paRNAs revealed 88 upregulated and 99 downregulated paRNAs in AD mFC regions (**Fig. 2E**, and **Supplementary Table 6**). These changes are also recapitulated by total RNA-seq data from Normal/AD brains by Nativio et al.^36^ (from LTL brain region, **Fig. 2F**). paRNAs altered in AD versus Normal brains are distinct from those changed in normal aging (old versus young human brains that are cognitively normal, **Fig. 2F**), suggesting potential involvement of these RNAs in AD pathogenesis. Interestingly, we found that paRNAs upregulated in AD are much longer than paRNAs downregulated in AD, a pattern not observed for deregulated protein-coding genes (**Supplementary Fig. 5G**). Although it is currently unclear what the molecular basis underlying this observation is, we speculate that there are malfunctions of RNA processing in AD that deregulate paRNAs in a length-related manner. Consistent with the correlation seen for mRNAs and other ncRNAs, hyper- and hypo-methylation of paRNA is correlated with their expression increase and decrease in AD, respectively (**Fig. 2G,H**).

### paRNA deregulation in AD mouse model and other human neurodegenerative diseases

We examined if the paRNAs deregulated in human AD brains can be detected in 5xFAD mice brain, a well-characterized mouse model of AD^45^. Total RNA-seq data has been generated from the right cerebral hemisphere of 5xFAD mice brain (ID: syn21983020). There are 1,378 paRNAs produced from promoters of homologous genes in human and mice brains (hereafter referred to as common paRNAs) (**Fig. 3A**). Of these, about 6% (n=87) showed altered expression in human AD brains (**Fig. 3B**), and about 3% (n=42) were differentially expressed in the 5xFAD mouse brains versus wild-type controls. However, none of them is consistent between human and mouse AD brains (**Fig. 3B**). Because 5xFAD is mainly an amyloid model, whereas our human RNA-seq data are from patients bearing both Tau and amyloid pathologies, we examined another mouse model exhibiting both these two pathologies, the 3xTg-AD model^46^. Based on the same process done above for 5xFAD, we identified 3,934 paRNAs in 3xTg-AD mouse brains (hippocampus, data from syn22964719). Differential expression analysis identified 5 upregulated and 1 downregulated paRNAs in 3xTg-AD mice (n=18) compared to wild-type (n=18). There are no common differentially expressed paRNAs between 3xTg-AD mouse model and human AD brains (**Fig. 3C**). A similar conclusion is also drawn by comparing paRNA changes in the human brain versus those changes in the P301S mouse brains^47^, a commonly used tauopathy model (**Fig. 3D**). Such a lack of similarity of paRNAs altered in human and mouse AD brains is consistent with the idea that AD mouse models cannot completely recapitulate human AD pathogenesis^48^.

**Figure 3.**
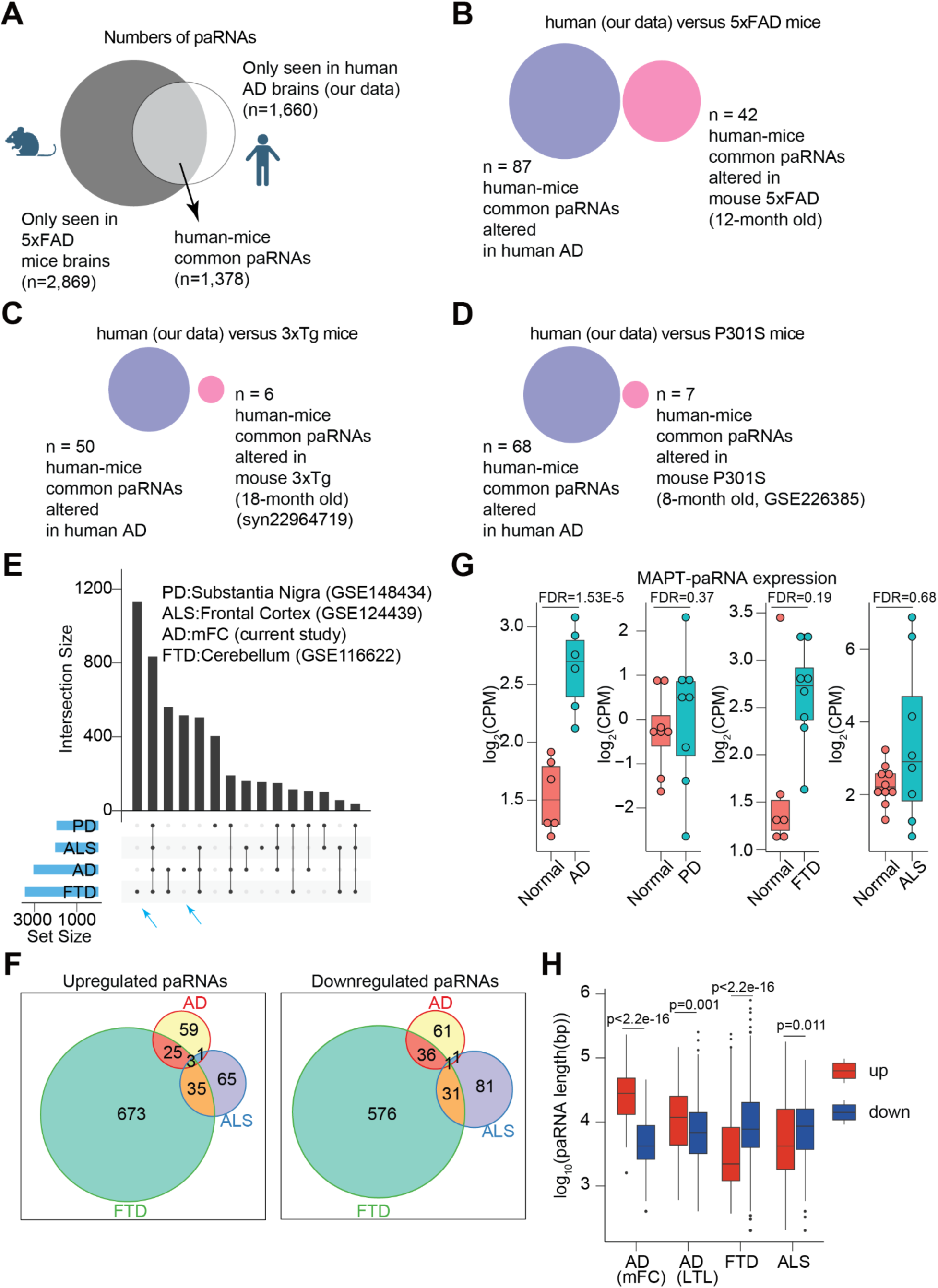
Comparing paRNA changes in human AD versus mouse AD models or other human neurodegenerative diseases. **A**. A venn diagram showing the numbers of human-mice common and species-specific paRNAs in human and mouse brains, respectively. The human data is from our current work. The mouse data is calculated from RNA-seq datasets generated in brains of a 5xFAD mouse model. Common paRNAs were defined by their production from promoters of homologous genes. **B-D**. Venn diagrams showing dysregulated paRNAs from human-mouse common paRNAs in either human brains or mouse models as indicated. **E**. An UpSet plot showing common and unique paRNAs detected in human brain RNA-seq data generated in donors bearing several neurodegenerative diseases. The datasets information and brain regions are indicated. Blue arrows point to paRNAs uniquely seen in FTD and AD. **F**. Venn diagrams showing the limited overlaps of upregulated and downregulated paRNAs across different neurodegenerative diseases, respectively. **G**. Boxplots showing the expression levels of *MAPT-paRNA* across different neurodegenerative diseases. FDR values were calculated by DESeq2. **H**. Boxplots showing the length of upregulated and downregulated paRNAs across different neurodegenerative diseases. Red and blue boxplots represent upregulated and downregulated paRNAs in each disease, respectively. P values were calculated by a two-tailed non-parametric Wilcoxon–Mann–Whitney test. In all panels, boxplots indicate the interquartile range with the central line representing the median, and the vertical lines extending to the extreme values in the group.

We next examined whether any paRNAs are also dysregulated in human brains with other neurodegenerative diseases, such as Parkinson’s disease (PD)^49^, Frontotemporal Dementia (FTD) ^50^, and Amyotrophic lateral sclerosis (ALS)^51^. For these, we retrieved RNA-seq datasets for each disease, and identified 1,826 paRNAs in PD, 3,454 in FTD, and 2,007 paRNAs in ALS, respectively, by *de novo* transcript calling (**Fig. 3E**). These paRNAs showed non-overlapping expression patterns in different diseases: for example, more than 500 and 1,000 paRNAs are only found in AD and FTD brain samples, respectively (blue arrows, **Fig. 3E**). We identified deregulated paRNAs in each disease, and these showed limited overlaps (**Fig. 3F**). These results demonstrate disease-specific expression or deregulation of paRNAs, although the contribution of RNA sample quality and different brain regions used in the data generation cannot be fully excluded. For example, *MAPT-paRNA* is expressed in several diseases, but it is upregulated only in the human AD brains (**Fig. 3G**, also see **Fig. 2E**). We observed that the length of upregulated paRNAs was longer than that of downregulated paRNAs in AD brains, which is reproducible by analyzing data from Nativio et al.^36^ (**Fig. 3H**). Interestingly, this pattern was not seen or even reversed in ALS and FTD (**Fig. 3H**). These results indicate that paRNAs can be deregulated in several neurodegenerative diseases, but the exact paRNAs and the mechanisms underlying their deregulation can be disease- or context-dependent.

### Deregulation of paRNAs in AD does not correlate with promoter epigenetic state changes

As paRNAs and their nearby genes share promoters, their expression levels are expected to be correlated if they are regulated at the transcriptional level by the promoters’ epigenetic states (**Supplementary Fig. 6A**). However, when we analyzed AD-associated expression changes of paRNAs versus the bidirectional mRNAs from the same promoters, we observed a poor correlation between upregulated paRNAs versus their bidirectional mRNAs (median correlation coefficient R=-0.01, n = 88 pairs); whereas the correlation between AD-downregulated paRNAs and their bidirectional mRNAs is better but remains modest (median correlation coefficient R=0.42, n = 99) (**Supplementary Fig. 6B**). Two representative examples are given, showing an AD-upregulated paRNA from the *CTCF* gene promoter, and an AD-downregulated paRNA antisense to the *VMA21* gene promoter (**Supplementary Fig. 6C**). This phenomenon was recapitulated in another independent AD dataset from the LTL region^36^ (**Supplementary Fig. 6D**). Taking *MAPT-paRNA* as an example, while it is consistently upregulated in AD, its neighboring gene *MAPT* was not significantly changed from either of two brain regions/cohorts we analyzed (**Supplementary Fig. 6E,F**). These findings suggest that AD-deregulated paRNAs, especially those upregulated in AD, are likely altered in a manner decoupled from the nearest mRNA genes; and if these paRNAs may have functions, it is unlikely they impact their promoter-sharing nearby genes.

We further examined the epigenetic states of deregulated paRNAs. Because we found consistent changes of paRNAs in the LTL regions of AD brains from the study by Nativio et al.^36^ (**Fig. 2F**), which generated ChIP-seq data for acetylation of histone H3 at residue K27 (H3K27ac), a histone marker for active promoters/enhancers, we thus tested this question in LTL. We found that the H3K27ac signal around TSSs of deregulated paRNAs was not different between Normal and AD brains (**Supplementary Fig. 6G**). This observation was recapitulated by analyzing ChIP-Seq datasets of H3K27ac and tri-methylation of histone H3 lysine 4 (H3K4me3) generated by a separate AD brain cohort (DLPFC regions, Rush Alzheimer’s Study) (**Supplementary Fig. 6H**). Taking *MAPT* and *TMEM41A* loci as examples, the H3K27ac and H3K4me3 signals on their promoters were similar between Normal and AD brains, despite these two paRNAs being upregulated (**Supplementary Fig. 6I**). In summary, our results indicate that the epigenetic state, at least by the histone markers we examined here at the TSSs, is not associated with paRNA deregulation in AD.

### Cell-type specific paRNA expression and deregulation in AD

Brain tissue consists of several neuronal and glial cell types. We reanalyzed a publicly available single-nucleus RNA-seq (snRNA-seq) dataset generated from Normal/AD human brains^52^ (DLPFC region) to appreciate expression patterns of paRNAs in different cell types. Different numbers of Unique Molecular Identifiers (UMIs) and cells of each major type were observed (**Supplementary Fig. 7A,B**). In total, 763 and 872 paRNAs we identified by bulk RNA-seq data of mFC can be detected by DLPFC snRNA-seq in Normal or AD conditions, respectively (**Supplementary Fig. 7C**), among which, m6A-marked paRNAs were more detectable (**Supplementary Fig. 7D,E**). These numbers were broken down to cell-type-specific patterns, and excitatory neurons (Ex) possess the largest numbers of paRNAs detected (**Supplementary Fig. 7F**). Among paRNAs seen in snRNA-seq data, about 30% (n=284) are exclusively detected in neurons (excitatory, Ex; or inhibitory, In), and a smaller group of paRNAs (n=69) are exclusively detected in glial cell types (such as astrocytes, Ast; oligodendrocytes, Oli; or microglia, Mic; **Supplementary Fig. 7F**). For the top five highly AD-upregulated paRNAs (**Fig. 2E**), they all displayed a certain degree of cell-type-specificity (**Supplementary Fig. 7G**). For example, *DOCK7-paRNA* was highly expressed in Ast; *MAPT-paRNA* is expressed higher in Ex and In, and to some degree in Oli (**Supplementary Fig. 7G**). By comparing Normal and AD conditions in each cell type, *MAPT-paRNA* was significantly increased in AD only in Ex (**Supplementary Fig. 7H**). This paRNA is annotated as *MAPT-AS1* in the Gencode database, but for consistency in this paper, we will refer to it as *MAPT-paRNA*. These results indicate that paRNAs are expressed in the human brain and altered in AD with cell-type-specificity.

### m6A methylation regulates stability of paRNAs

We searched for appropriate cell models to study the potential roles of AD-associated paRNAs and the impact of m6A. In major iPSC-derived brain cells where RNA-seq data are available^53^, each of the top five AD-deregulated paRNAs was found to display some level of cell-type-specificity, but they are more often expressed highly in neurons (**Supplementary Fig. 8A**). Among these, *MAPT-paRNA* exhibited the highest and almost exclusive expression in iPSC-derived neurons (**Supplementary Fig. 8A**). It is notable that the cell-type-specificity of paRNAs revealed by DPFLC snRNA-seq data differs from that by bulk RNA-seq from iPSC derived cells (**Supplementary Figs. 7G, 8A**), indicating that cell models can only partially recapitulate paRNA expression patterns in adult and/or AD human brains.

*MAPT* is a central disease locus in AD, because it encodes for the Tau protein that drives neurofibrillary tangles in AD brains^3^. A previous study reported that *MAPT* nature antisense transcript (NAT), which largely overlaps and is essentially *MAPT-paRNA*, represses *MAPT* translation to Tau protein in SH-SY5Y cell line^33^. However, another study found that *MAPT-paRNA* does not impact *MAPT*/Tau at either the transcriptional or translational levels^34^. Consequently, the functional role of *MAPT-paRNA* in AD remains unclear. We thus elected to focus on *MAPT-paRNA* for in depth experimental studies. We first tested the expression of *MAPT-paRNA* in cellular models, including human embryonic stem cells (H1-ESCs), a human microglia cell line (HMC3), a human neuroblastoma cell line (SH-SY5Y), H1-hESC-derived astrocytes, neural progenitor cells (NPC), and iPSC-derived microglia and neurons (i3Neurons) (see methods, **Fig. 4A**). Quantitative reverse-transcription PCR (qRT-PCR) results confirmed that *MAPT-paRNA* is highly and almost exclusively expressed in i3Neurons as compared to other cell types (**Fig. 4A**). The i3Neurons we generated are based on a doxycycline (Dox) inducible NGN2-directed differentiation method of WTC11 iPSCs^54^, which are glutamatergic excitatory neurons that displayed well-recognized neuronal maturation processes by both morphology and molecular markers (**Supplementary Fig. 8B,C**). Three weeks after in vitro differentiation (IVD), these neurons started to express *MAP2*, a neuronal marker, and by 6-8 weeks, they showed prominent expression of synaptic markers, such as postsynaptic density protein 95 (PSD95) (**Supplementary Fig. 8B,C**). *MAPT-paRNA* appeared to gradually increase expression during neuronal maturation and its level plateaued at around 5 weeks IVD (**Supplementary Fig. 8D**). To investigate how m6A methylation may impact *MAPT-paRNA* expression, we treated 8-week-old i3Neurons with STM2457^55^, a chemical inhibitor of m6A writer METTL3. Antibody based m6A RIP-qPCR showed that this treatment reduced the m6A methylation level of *MAPT-paRNA*, which resulted in its expression decrease as shown by RT-qPCR (**Fig. 4B,C**). These results suggest that m6A positively regulates *MAPT-paRNA* expression. In contrast, this treatment did not impact either the mRNA expression or transcription level of *MAPT*, which shares the same promoter with the paRNA. These are revealed by using primers targeting the mRNA (across exons) or the intronic regions (indicating pre-mRNA transcription), respectively (**Supplementary Fig. 8E**). These results indicate that m6A inhibition did not reduce *MAPT-paRNA* level via altering the transcriptional activity of the shared promoter, which otherwise would have affected *MAPT* transcription as well. We further examined if the observed *MAPT-paRNA* reduction takes place at the RNA stability level, and found that after transcriptional inhibition, the paRNA with a lower m6A level (i.e., presence of STM2457) displayed a lower stability and faster decay (**Fig. 4D**). We additionally conducted genetic knockdown (KD) of METTL3 by an antisense oligonucleotide (ASO) in 8-week old i3Neurons (**Supplementary Fig. 8F,G**), which reduced both m6A modification and RNA expression level of *MAPT-paRNA* (**Supplementary Fig. 8H,I**), consistent with the results from the chemical inhibitor treatment.

**Figure 4.**
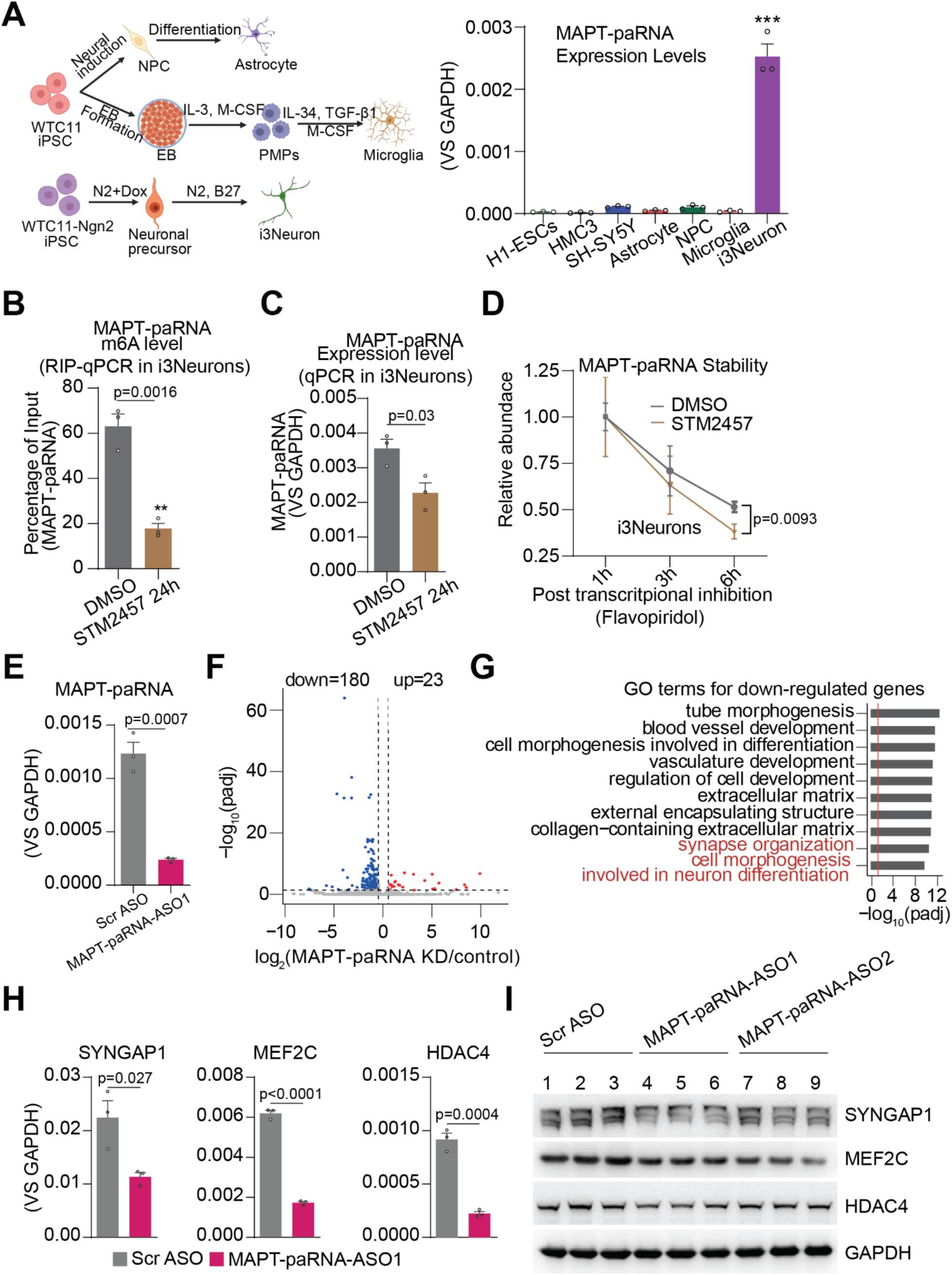
Neuron-enriched *MAPT-paRNA* stability control by m6A and its global regulatory functions. **A.** (Left) Cartoon diagrams illustrating the differentiation of iPSC-derived brain cells (created with BioRender.com). (Right) A barplot showing the expression levels of *MAPT-paRNA* in various cell types. P value was calculated by a two-tailed student’s t-test. **B,C**. MeRIP-qPCR and RT-qPCR data showing m6A methylation and *MAPT-paRNA* expression with and without STM2457 treatment, respectively. **D**. Time course stability of *MAPT-paRNA* measured by RT-qPCR after transcriptional inhibition, with and without STM2457 pre-treatment for 24hr. P value was calculated by a two-tailed student’s t-test. **E**. A barplot showing the relative RNA expression of *MAPT-paRNA* quantified by RT–qPCR after scramble or targeting ASO treatment. **F**. A volcano plot showing differentially expressed genes after *MAPT-paRNA* knockdown using a targeting ASO versus the scramble ASO treatment. Red and blue dots represent upregulated and downregulated genes, respectively. **G**. GO analysis for the downregulated genes after *MAPT-paRNA* knockdown. The red line represents adjusted p values (padj) at 0.05. **H**. Barplots showing relative RNA expression of *MEF2C*, *SYNGAP1*, and *HDAC4* quantified by RT–qPCR. P values were calculated by a two-tailed student’s t-test. **I**. Protein levels of *MEF2C*, *SYNGAP1*, and *HDAC4* showing triplicates of Western blotting (WB) after *MAPT-paRNA* knockdown.

### *MAPT-paRNA* regulates global gene transcription in human neurons *in cis* and *in trans*

We further studied the functions of *MAPT-paRNA* by employing ASOs to KD it in 8-week IVD excitatory i3Neurons, because at this stage, neurons possess maximal level of expression of this paRNA, displaying prominent neuronal morphology and synaptic features (**Supplementary Fig. 8B,C**). Gymnotic delivery of ASO achieved potent KD of *MAPT-paRNA* (>80% reduction) without affecting neuronal viability or morphology (**Fig. 4E, and Supplementary Fig. 9A**). We found that KD of *MAPT-paRNA* did not alter *MAPT*/Tau at either the transcriptional or translational levels in i3Neurons (no matter if it is the mRNA, the Tau protein, or the phosphorylated Tau (p-Tau) that is implicated in AD pathology), even after an extended 7-day KD (**Supplementary Fig. 9B,C**). This indicates that *MAPT-paRNA* did not regulate the neighboring gene, *MAPT*, consistent with the report by Policarpo, et al.^34^. No change of *MAPT*/Tau was seen after the KD of *MAPT-paRNA* in SH-SY5Y cells, either (a neuroblastoma cell model used by a prior study of MAPT-paRNA^33^) (**Supplementary Fig. 9D,E**). This result is consistent with the fact that paRNAs upregulated in AD show poor correlation with changes of their bidirectional mRNAs (**Supplementary Fig. 6A,B,D,E,F**).

To unbiasedly explore its functions, we performed RNA-seq after *MAPT-paRNA* KD. This revealed over 200 genes deregulated, with most of them showing downregulation (n=180), and a smaller subset upregulation (n=23) (**Fig. 4F**, and **Supplementary Table 8**). RNA-seq also confirmed qPCR data that *MAPT* mRNA was not changed (**Supplementary Figs.9B,10A**). GO analysis revealed that the functions of these downregulated genes are closely related to synaptic functions (**Fig. 4G**, and **Supplementary Table 9**). Among these, some are known to be important regulators of neuronal/synaptic biology, such as *SYNGAP1*^56^*, MEF2C*^57^, and *HDAC4*^58^ (**Supplementary Fig. 10B,C,D**). We validated the expression reduction of these “target genes’’ of *MAPT-paRNA* by RT-qPCR (**Fig. 4H**) and by a second ASO (**Supplementary Fig. 10E**). Consistently, the protein levels encoded by these target genes were also decreased (**Fig. 4I**, and **Supplementary Fig. 10F**). These results demonstrated that *MAPT-paRNA* is a global regulator of gene expression that selectively impacts neuronal/synaptic functions. Interestingly, very few of these target genes are located in its genomic vicinity, and most of them are on different chromosomes (**Supplementary Fig. 10G**), suggesting that *MAPT-paRNA* regulates target gene expression both *in cis* and *in trans*. This is a distinct function from the reported action of some paRNAs that impacts the bidirectional gene sharing promoter with them^27,28^.

### Functional paRNA-target-gene interactions in 3D nucleus revealed by single-cell RNA-DNA interactome

A plausible mechanism underlying paRNAs’ roles in gene expression control is that they may directly interact with target genes via spatial proximity in a 3D nucleus. To explore this, we analyzed recent single nuclei RNA-DNA interactome data of the human brain (DLPFC region), namely by the Multi-Nucleic Acid Interaction Mapping in Single Cell (MUSIC) technique^59^ (**Fig. 5A**). MUSIC employs cell barcodes (CB) to define molecules (DNA or RNA) from the same cell, and then uses molecular barcodes (MB) to identify shared molecular complexes that define interactions of DNAs or RNAs (**Fig. 5A**, **see methods**). It provides single cell RNA-DNA interactome and thus can reveal paRNAs’ cell-type specific target genes. The cell numbers and gene expression counts from our analysis of the MUSIC datasets are shown in **Supplementary Fig. 11A,B** and are identical to what was originally reported^59^.

**Figure 5.**
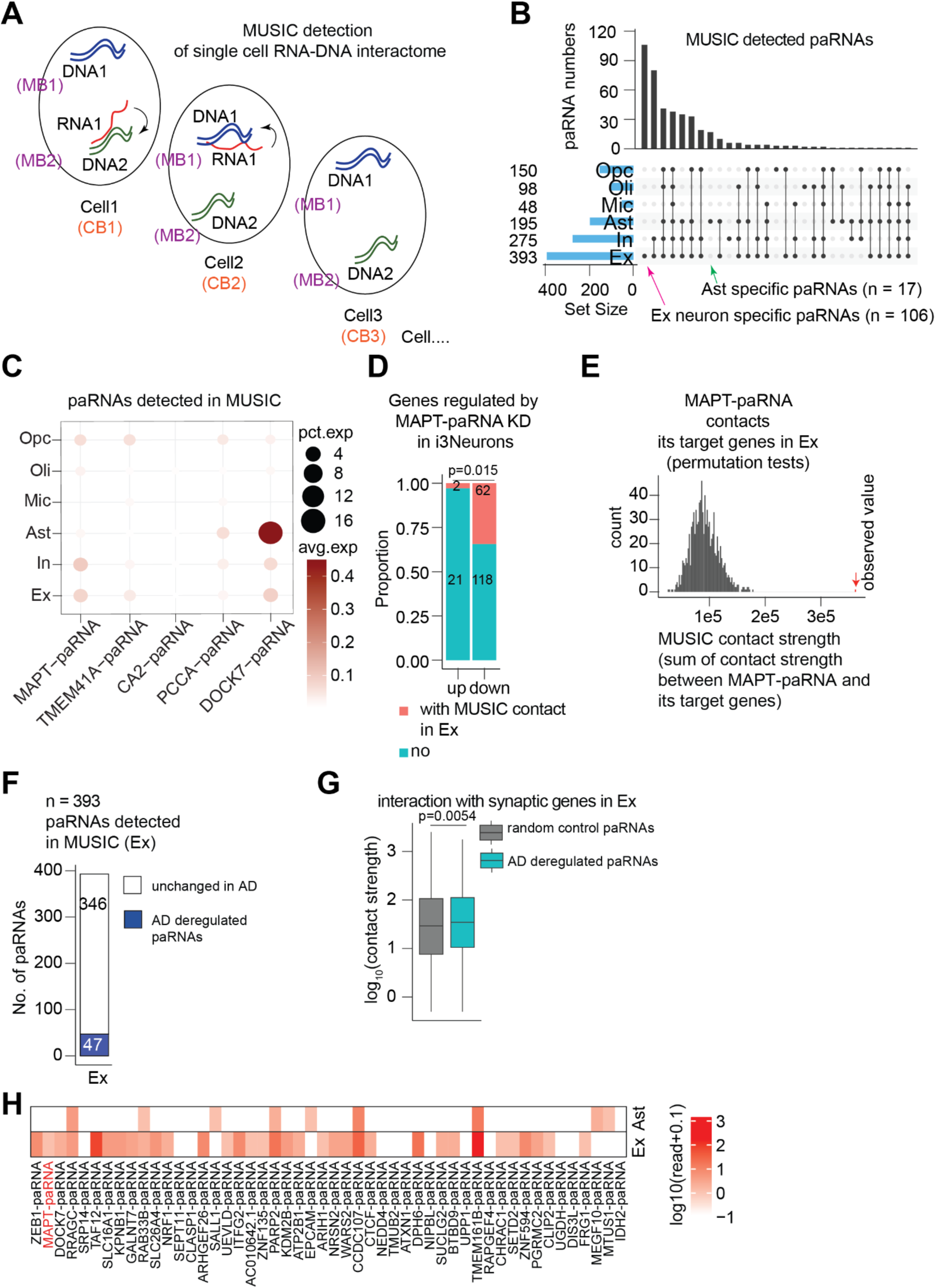
paRNA-DNA interaction in human brain cells revealed by MUSIC data. **A**. A diagram showing RNA-DNA interaction at single-cell level in MUSIC data using cell barcodes (CB) and molecular barcodes (MB). CB is shared by all molecules in one cell, and MB is shared by all molecules in one molecular complex. **B.** An UpSet plot showing paRNAs and their numbers (to the left) in different cell types detected by MUSIC from Normal brain. **C**. A dotplot showing the average expression of the top five AD-upregulated paRNAs in MUSIC data. Ex: Excitatory neurons. In: Inhibitory neurons. Ast: Astrocyte. Mic: Microglia. Oli: Oligodendrocyte. Opc: Oligodendrocyte progenitor cell. **D**. A barplot showing the numbers of differentially expressed genes seen after MAPT-paRNA KD that form RNA-DNA interactions with *MAPT-paRNA* in the Ex neurons. P value: Fisher’s exact test. **E**. A barplot from permutation test showing the observed contact strength between MAPT-paRNA and its true 62 target genes (red arrow to the right) versus contact strength between MAPT-paRNA and 1000 sets of randomly selected 62 genes. **F**. A barplot showing the number of paRNAs detected by MUSIC data in Ex neurons and the subset that show AD-deregulation in bulk RNA-seq data in Fig.2. **G**. A boxplot showing the RNA-DNA contact strength between the two groups of paRNAs and synaptic genes in Ex: AD-deregulated paRNAs (n = 47 in panel F), and non-AD-deregulated paRNAs (n = 346). P value: two-tailed non-parametric Wilcoxon–Mann–Whitney test. Boxplots indicate the interquartile range with the central line representing the median, and the vertical lines extending to the extreme values in the group. **H**. A heatmap showing the RNA-DNA contact strength between AD-deregulated paRNAs and a single *MEF2C* gene detected in Ex and Ast, respectively.

Overall, 425 paRNAs can be detected in MUSIC data in at least one of the cell types. Among these, Ex exhibited the largest number of detectable paRNAs (n = 393, **Fig. 5B**). This is perhaps partially because our paRNA list was identified in bulk brain RNA-seq data in which neuronal signals are dominant ^38^, and partially it may also be due to higher detectable RNA reads in Ex in MUSIC data (**Supplementary Fig. 11A,B**). For each paRNA, MUSIC identified their interacting genes in the 3D nucleome based on shared CBs and MBs (**Fig. 5A, see methods**). Ex and Ast exhibited 106 and 17 paRNAs that are specific/exclusive to these two cell types, respectively (**Fig. 5B**). We identified genes showing RNA-DNA contacts with these cell-type-specific paRNAs in their respective cell types, and found 4,736 interacting genes for Ex-specific paRNAs and 1,055 genes for Ast-specific paRNAs. GO analysis of these paRNA-interacting genes revealed them to be related to neuron differentiation/synapse in Ex and cell-cell adhesion in Ast (**Supplementary Fig. 11C**), respectively, suggesting that cell-type-specific paRNAs may play roles in regulating cell-type-specific genes and cellular functions.

We found that m6A-paRNAs can be better detected by the MUSIC data than non-m6A-modified paRNAs, and this is consistent in all cell types (**Supplementary Fig. 11D,E**). This pattern is similar to that in snRNA-seq data (**Supplementary Fig. 7D,E**). As compared to non-m6A-paRNAs, m6A-paRNAs on average can interact with more genes, and show higher interaction strength with genes they contact, a trend particularly obvious in Ex (**Supplementary Fig. 11F**). These results support a notion that m6A-paRNAs are more detectable in MUSIC data possibly due to higher stability, and they possess higher propensity to act in target gene regulation, particularly in Ex. The top five AD-upregulated paRNAs display expression in two or three cell types in MUSIC data (**Fig. 5C**), again a pattern similar to that revealed by snRNA-seq (**Supplementary Fig. 7G**). As aforementioned, KD of *MAPT-paRNA* in i3Neurons, a model of human excitatory neurons, deregulated ∼200 genes (**Fig. 4F,G**), many of which are important for neuron differentiation and synapse organization. To test the mechanisms, we examined if *MAPT-paRNA* forms direct RNA-DNA contacts with these “target genes”, and we found that about 30% of them (up=2 and down=62) indeed displayed MUSIC RNA-DNA contacts in Ex (**Fig. 5D**). Given that only 2 upregulated genes interact with *MAPT-paRNA*, we will focus on the 62 downregulated “target genes” interacting with *MAPT-paRNA*. The observed contact strength between *MAPT-paRNA* and these 62 target genes was much higher than by random chance (**Fig. 5E**), as shown by permutation tests, for which we randomly selected 62 genes 1,000 times, and calculated the contact strength between *MAPT-paRNA* and each of these random genesets. The contacts between *MAPT-paRNA* and the 62 target genes were more pronounced in Ex than in In or Ast (**Supplementary Fig. 11G**), lending support to the notion that *MAPT-paRNA* contacts these genes and functionally activates their expression specifically in Ex.

We attempted to test whether this notion may apply to other paRNAs in Ex; i.e., do many of them contact synapse/neuronal genes to potentially play roles in gene regulation and in AD? Out of 393 detectable paRNAs in Ex in MUSIC data, 47 were deregulated in our bulk RNA-seq data (**Fig. 5F**). We calculated RNA-DNA contact strength of each of these paRNAs with a curated set of synapse-associated genes (n = 393, see methods about this curated list), and we found that paRNAs deregulated in AD, as compared to average paRNAs, display higher interaction frequency with synapse-associated genes in Ex (**Fig. 5G**). As an example, RNA-DNA contacts were found in Ex to form between not only *MAPT-paRNA,* but also a series of other AD-deregulated paRNAs, with *MEF2C* (encoding a key neuronal factor known to confer resilience to AD^47,60^)(**Fig. 5H**). In contrast, these interactions did not take place in Ast (**Fig. 5H**). These results support that at least a subset of neuronal paRNAs, as exemplified by *MAPT-paRNA*, can contact and regulate neuronal/synapse genes; alteration of such paRNAs in AD are involved in deregulation of key synaptic genes and AD pathogenesis.

### *MAPT-paRNA* protects neurons from excitotoxicity by modulating synaptic activity

An important feature of AD is neuronal hyperexcitability, which leads to excitotoxicity and neuronal death^61,62^. This is often mediated by an overactive N-methyl-d-aspartate receptor (NMDAR) at the synapse. Several key synaptic genes dependent on *MAPT-paRNA*, such as *MEF2C* and *SYNGAP1* (**Fig. 4G,H**), have been known to modulate NMDA activity at synapses ^63–65^. Indeed, *Mef2c* deletion in mice impairs learning and memory^66^, and it has been shown to prevent neuronal apoptosis elicited by excitotoxic NMDA^65^. Loss of *SYNGAP1* induces neuronal hyperexcitability, leading to deficits in learning and memory^67^. These results suggest that *MAPT-paRNA* might act as a neuroprotective modulator in the context of AD.

To test this, we employed an AD-relevant glutamate-induced excitotoxicity model^68,69^ (**Fig. 6A**). We found that mature i3Neurons (8-week IVD) with *MAPT-paRNA* KD are more vulnerable to glutamate-induced cell apoptosis, as shown by significantly higher numbers of cells showing Propidium Iodide (PI) staining (**Fig. 6B,C**). Since SYNGAP1 and MEF2C have been reported to inhibit excitotoxicity by modulating the levels of postsynaptic proteins and glutamate receptors, we selected four of these proteins/receptors to test, finding PSD95 and GluN1 (glutamate NMDA Receptor 1) significantly increased in i3Neurons after *MAPT-paRNA* knockdown, while Synapsin1 (a presynaptic protein) and GluA2 (glutamate AMPA Receptor subunit 2) were not changed (**Fig. 6D**, and **Supplementary Fig. 12A**). These changes did not take place at the mRNA levels (**Supplementary Fig. 12B,C**), consistent with the reports that SYNGAP1 or MEF2C modulate post-synapse proteins and excitability locally at the synapse^70,71^. We also conducted KD of *MEF2C* and *SYNGAP1* mRNAs by ASOs, and these caused upregulation of PSD95 and GluN1, but not GluA2, a pattern consistent with *MAPT-paRNA* KD (**Supplementary Fig. 12D,E,F**). Furthermore, immunostaining after *MAPT-paRNA* KD directly showed accumulation of PSD95 and GluN1 at the dendrites of i3Neurons (**Fig. 6E,F**). Importantly, neuronal electrophysiology confirmed alteration of NMDA receptor activity after KD of this paRNA, while AMPA receptor activity was unaffected (**Fig. 6G,H**). The protective role of *MAPT-paRNA* also applies to beta-amyloid (Aβ-42)-induced neuronal cytotoxicity^72,73^, as revealed by increased lactate dehydrogenase (LDH) release in the culture media if i3Neurons were depleted of this paRNA (**Supplementary Fig. 12G**). All together, these results indicate that *MAPT-paRNA* exerts neuroprotective effects, at least in part, via regulating the expression of key synapse-associated genes to modulate the distribution and assembly of synaptic scaffolding proteins and glutamate receptors (**Fig. 6I**).

**Figure 6.**
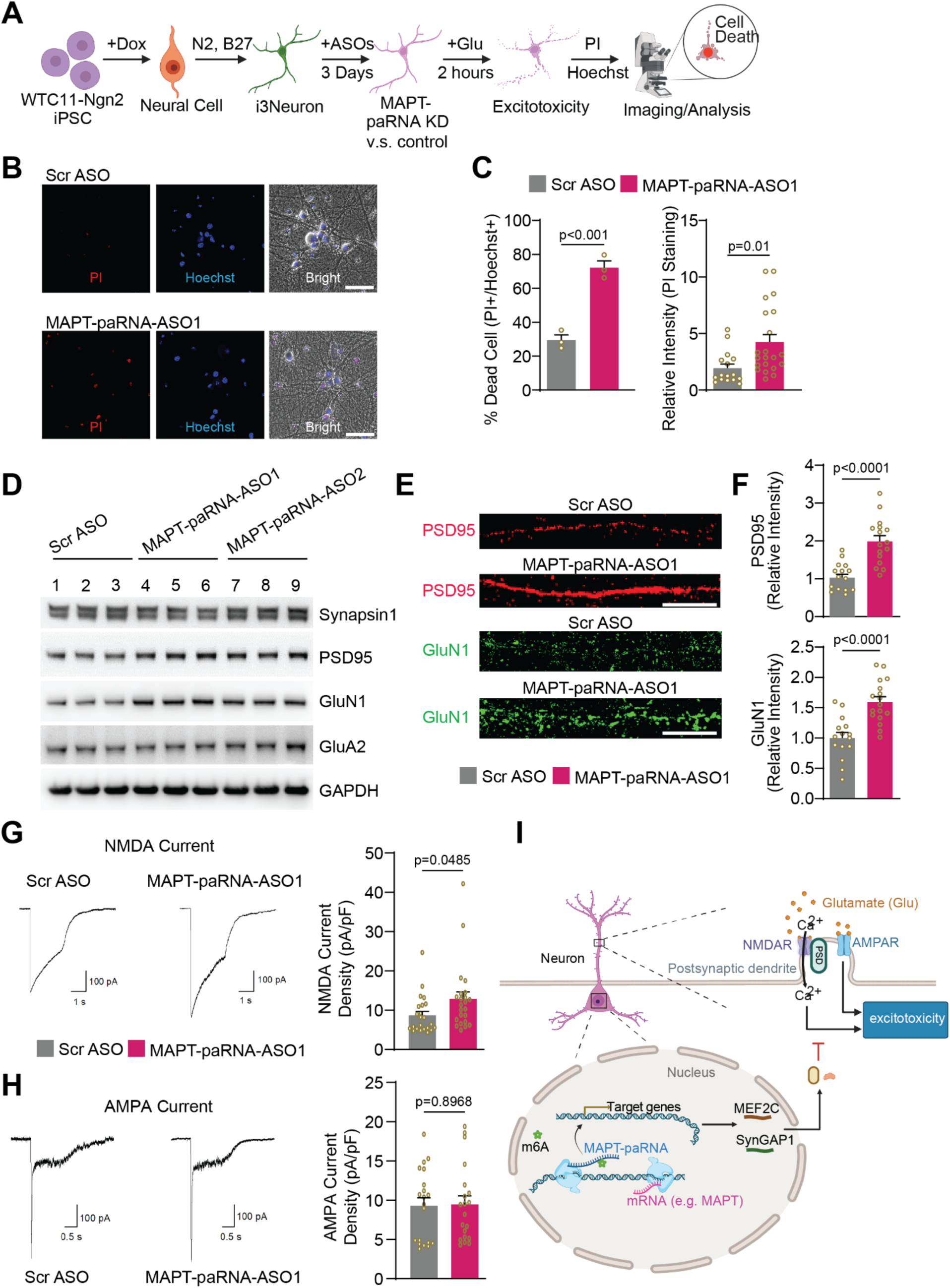
The neuroprotective role of *MATP-paRNA* against excitotoxicity. **A**. Schematic representation of glutamate toxicity in WTC11-derived i3Neurons. 8-week-old i3Neurons with/without *MAPT-paRNA* knockdown were treated with 10µM glutamate for 2 hours. PI (propidium iodide) and Hoechst 33342 were introduced to the medium to visualize dead (PI+) and total (Hochest+) cells, respectively. **B.** Representative images showing glutamate-induced i3Neuron cell death after *MAPT-paRNA* knockdown. Scale bar, 50µm. **C**. Quantification of apoptotic i3Neurons from panel B. **D**. Triplicates of western blot analysis of synaptic markers (Synapsin1 and PSD95) and glutamate receptors (GLUN1 and GLUA2) in i3Neurons after *MAPT-paRNA* knockdown. **E**. Representative images showing immunofluorescent signals of PSD95 and GluN1 at the dendrites of i3Neurons after *MAPT-paRNA* knockdown, showing their accumulation. Scale bar, 10µm. **F.** Quantitative analysis of signals in panel **E**. For each group, at least 15 neurons from 3 different experiments were quantified and analyzed. **G.** Representative NMDA current traces (left panel) and current density comparison (right panel) from i3Neurons treated with scramble control and *MAPT-paRNA* ASO1 (n = 23 neurons per group). **H**. Representative AMPA current traces (left panel) and current density comparison (right panel) from i3Neurons treated with scramble control and *MAPT-paRNA* ASO1 (n = 20 neurons per group). **I**. A model of *MAPT-paRNA* function, as a representative m6A-modified ncRNA in AD, in neuronal gene regulation and survival (generated by Biorender). P values: two-tailed unpaired student’s t-tests were used for **C, F**, **G**, and **H**.

## Discussion

### m6A methylome on coding and noncoding RNAs in human brain and AD

Our current study utilized total RNA-seq and m6A RIP-seq datasets in human brain samples from AD patients or cognitively normal controls, and identified a comprehensive landscape of RNA m6A methylome on mRNAs and ncRNAs. Our results showed that about 60-70% of m6A peaks in the human mFC are located in the noncoding regions (**Fig.1A**, and **Supplementary Fig. 2H**), encompassing various categories of ncRNAs. These include annotated lncRNAs or other ncRNAs (e.g., snoRNAs and snRNAs), as well as ncRNAs that have not been previously annotated in the human transcriptome (Gencode database), including many paRNAs that we focused on in the latter part of this work. Interesting, we observed around 8,000 highly m6A-marked intronic regions that overlap with L1 elements. These intronic m6A-marked L1s have been identified by our previous work in cancer cells, fetal human brains, and iPSC-derived neural progenitor cells^12^, in which we dubbed them m6A-marked intronic L1s (MILs) and found they can impact the transcription of the genes that harbor them. While in this current work we focus on paRNAs for functional studies because they have been less explored, it will be interesting and important to further test the roles of various m6A-modified ncRNAs in gene regulation, cellular functions in the human brain, and in AD pathogenesis. Therefore, our current work offers an m6A methylome that has significantly extended the previous observations of m6A on mRNAs in normal human brains or in diseased brains from AD or ALS^19,74^.

Our RNA-seq transcriptome data captured known features of the AD brain, including reduced expression of synapse-related genes, and this pattern is consistent with other recent datasets^36^. Interestingly, the changes of m6A methylome in normal and AD conditions are not particularly pervasive, i.e., we identified 835 and 1,645 regions showing m6A hypo- and hyper-methylation. This selective and moderate m6A change in AD is consistent with the fact that we did not observe strongly changed gene expression of m6A modifying enzyme genes in AD brains, such as *METTL3* and *METTL14.* In addition, we observed a moderate but significant positive correlation between m6A changes and ncRNA expression changes (**Supplementary Fig. 4A,B**).

This indicates that collectively m6A modification may play a positive role in regulating ncRNA expression in human brains. However, it is noteworthy that m6A is by no means the only mechanism that impacts the expression of these ncRNAs in the brain or in AD, and thus it is not surprising that the positive correlation is not particularly strong. Knockdown of METTL3, the main m6A enzyme, quantitatively affected *MAPT-paRNA* expression (**Supplementary Fig. 8F,G,I**). Other mechanisms affecting ncRNA expression in AD can be from the levels of transcriptional/epigenetic control and other non-m6A-mediated RNA stability regulation.

### Expression patterns and regulation of paRNAs in neurodegeneration

We identified 3,038 paRNAs in the human brain mFC region via *de novo* transcript calling, and found that close to 40% of them harbor m6A methylation. Interestingly, the genes neighboring m6A-marked paRNAs show enrichment of terms like synapses, indicating that paRNAs are produced and methylated selectively from important synapse/brain associated loci. By comparing paRNA landscapes we detected in human AD brain with those from mouse AD models, or from brain samples with other neurodegenerative diseases (NDDs), our study supports a conclusion that human AD-associated paRNAs are quite unique to human AD, and not commonly seen changed in other NDDs that we analyzed. These results demonstrate that paRNAs are potentially disease specific ncRNAs that play roles in gene dysregulation or disease etiology of NDDs.

Our analysis of available epigenetic datasets supports that AD-deregulated paRNAs, particularly those that show upregulation in AD, are unlikely to be altered at the epigenetic levels, as H3K4me3 and H3K27ac signals at their transcriptional start sites are similar in Normal and AD conditions. The expression changes of paRNAs and paired promoter-sharing mRNAs displayed poor correlation, too. This argues against that transcriptional/epigenetic deregulation is a main reason for paRNA alteration in AD, as this otherwise should deregulate the promoter-sharing gene mRNAs in a correlated manner. These results suggest a possibility that paRNA deregulation in AD may happen at the post-transcriptional level, likely as a consequence of defective RNA processing. Consistent with this, chemical inhibition of m6A reduces the stability of *MAPT-paRNA* from a key AD locus (i.e., *MAPT*), lending support that m6A methylation acts as at least one of the mechanisms in AD that can impact paRNA expression via stability control.

### *MAPT-paRNA* does not regulate neighboring *MAPT* mRNA or Tau protein levels

Given the central role of Tau protein in the pathology of AD and other neurodegenerative diseases, the locus coding Tau, *MAPT*, has been actively investigated to understand its mRNA expression and translational control^75^. AD-associated genetic variants exist in this locus^76,77^, and ncRNAs are produced therein. It is important to discover AD-associated ncRNAs that can regulate *MAPT*/Tau expression, which may offer mechanistic insights into Tauopathy in AD and potentially new therapeutic avenues. Simone et al. reported that <u>n</u>atural <u>a</u>ntisense transcripts (named NAT in their work), which is essentially *MAPT-paRNA*, can play a role to suppress *MAPT* mRNA translation^33^, but this result was not agreed upon by another study^34^. In our work, we found that *MAPT-paRNA* displayed much higher expression in iPSC-derived i3Neurons than cell lines or other types of iPSC-derived brain cells, and it reached a stable expression in morphologically mature excitatory neurons (i.e., with more than 5 weeks *in vitro* differentiation). We considered i3Neurons to be a more relevant model for studying functions of this and other brain paRNAs. Our knockdown of *MAPT-paRNA* did not impact *MAPT* expression either at transcriptional or protein levels. This result was also consistent in SH-5YSY, a neuroblastoma cell line model that displays low levels of *MAPT-paRNA*. In addition to the knockdown results, the poor correlation between paRNA/mRNA changes in AD also supports that if paRNAs bear functional roles, they unlikely regulate the genes directly adjacent to them. Furthermore, during m6A/METTL3 inhibitor treatment (STM2457), while *MAPT-paRNA* expression was reduced, the neighboring *MAPT* mRNA did not change. Together, our data are in agreement with a conclusion that *MAPT-paRNA* does not regulate the neighboring *MAPT* expression or translation.

### *MAPT-paRNA* acts as a global regulator of neuronal/synaptic gene expression and protects neuronal survival

Instead of regulating the neighboring *MAPT* gene, we found that *MAPT-paRNA* knockdown in mature i3Neurons altered around 200 genes, including both genes on the same chromosome and a large majority on other chromosomes. This indicates that *MAPT-paRNA* plays a regulatory role both *in cis* and *in trans*. By integrative analysis of the target genes and RNA-DNA interactome data in the human brain offered by the MUSIC dataset^59^, we found that the function of *MAPT-paRNA* can be explained by its direct interaction with target genes in the 3D nucleus of excitatory neurons. Interestingly, from MUSIC as well as snRNA-seq data, m6A-modified paRNAs are more detectable. Also, m6A-modified paRNAs tend to display more target genes and stronger contact strength with their target genes. These together support a positive correlation between m6A and paRNA stability/functionality. In addition to *MAPT-paRNA*, our analysis also found that cell-type-specific paRNAs in neurons often contact important neuronal/synaptic genes. Many AD-deregulated paRNAs in Ex neurons can contact synaptic genes more than background levels. These results indicate that many paRNAs may play roles in cell-type-specific gene regulation in human brains. Distinct from the past work that often found expression of paRNAs positively correlates with^23–26^, and in some cases functionally impacts, the expression of the neighboring gene sharing a promoter^27,28^, our current findings offer a renewed understanding of paRNAs as global regulators of gene expression *in cis* and *in trans*.

It is interesting that despite that many paRNAs can contact the same DNA loci (e.g., *MEF2C*), knockdown of a single *MAPT-paRNA* can already cause obvious change of *MEF2C* transcription. We found that in MUSIC data these paRNAs mostly engage with the target DNA alone, i.e., there are rarely any molecular complexes that contain more than one paRNA (data not shown). It is unknown how many of the 31 paRNAs that interact with *MEF2C* loci in Fig. 5H can play a functional role in augmenting *MEF2C* transcription - it could be a few, and some could even suppress *MEF2C* transcription. It is noteworthy that *MAPT-paRNA* may also impact the transcription of these other paRNAs, which we noticed in i3Neurons with *MAPT-paRNA* knockdown. We speculate that the *MAPT-paRNA* inhibition results in a collateral damage to a putative “paRNA network” that could reduce the transcription of *MEF2C* gene from two layers: 1), directly from the loss of *MAPT-paRNA*; 2) indirectly from other paRNAs that are affected by *MAPT-paRNA* loss. RNA transcription is being increasingly appreciated to take place in bursts^78^. We propose that many paRNAs are produced in dynamic transcriptional bursts, and they act via dynamic interaction with target gene DNAs to achieve transcriptional regulation, potentially in a collaborative manner.

As *MAPT-paRNA* showed increased expression in AD, we further explored its relevance to AD disease etiology. A glutamate-induced neuronal excitotoxicity assay showed that *MAPT-paRNA* plays a protective role under these conditions, which can be at least partially explained by its positive regulation of key synaptic regulators such as MEF2C and SYNGAP1. Similar protective role of this paRNA was found when neurons were challenged with Aβ-42 directly, one of the most known neuronal stresses in AD. We are tempted to speculate that *MAPT-paRNA* and perhaps some other regulatory paRNAs are upregulated in AD brains to safeguard key synapse/neuronal gene expression so as to rectify the declining neuronal functions. However, the full mechanisms underlying such paRNA regulation in human AD brains remain to be determined at this stage. Future research is warranted to dissect the cause of paRNA deregulation in AD, as well as the detailed mechanisms as to how individual paRNAs can regulate target genes in a 3D nucleus.

## Limitations of the current study

We utilized six brain samples from NCI and AD donors in this study, which in our knowledge, represented one of the largest that studied total RNA m6A methylome in human brains from normal/AD. However, this number remains limited, and some of the differential m6A sites revealed by current work will need additional larger datasets to be fully validated. Future work may also aim to profile RNA m6A methylome from multiple brain regions and/or from purified individual brain cell types to achieve higher temporal and spatial resolution. In this work we employed the MUSIC data to study RNA-DNA interactomes of paRNAs to understand the mechanisms of their action in gene regulation, but we observed that the detectability of paRNAs in MUSIC^59^ is lower than that of snRNA-seq data^52^. For example, about 10-20% of Ex neurons in snRNA-seq can detect *MAPT-paRNA* expression but this number ranges from one to a few percent in MUSIC. Therefore, the target gene numbers and DNA contact strengths of paRNAs seen in MUSIC may represent an under-estimation, and the paRNA-DNA interactome changes in AD cannot be confidently calculated at this stage. In addition, regarding the role of m6A, we noted that the change of *MAPT-paRNA* expression appeared to be quantitative after m6A inhibition (by either chemical or genetic approaches). This suggests that m6A is likely one of the mechanisms, rather than the only, that can impact *MAPT-paRNA* expression in human brains.

In summary, our results provided a blueprint of the m6A methylome in human brain and AD disease, particularly on various ncRNAs. We reveal previously unknown roles of paRNAs, as exemplified by *MAPT-paRNA,* as global transcriptional regulators in human neurons via navigating 3D nuclear organization, offering new insights into the epitranscriptome regulation of brain gene expression, neuronal survival, and AD pathogenesis.

## Methods

### Human brain sample collection

Together with Dr. Hui Zheng (Baylor College of Medicine), we collected RNAs from the middle frontal cortex (FC) of 12 Normal and AD patients (**Supplementary Table 1**). Paired total RNA-seq and m6A RNA methylome (m6A RIP-seq) data were generated from these samples. These tissues were originally collected by the Center for Neurodegenerative Disease Research (CNDR) from University of Pennsylvania and were provided to Dr. Hui Zheng.

### Cell culture

H1-hESCs (WA01) and WTC11 iPSCs were purchased from WiCell and Coriell, respectively. WTC11-Ngn2 iPSCs with doxycycline-inducible Ngn2 expression cassette was generously provided by Drs. Li Gan^54^ and Yin Shen (UCSF). Both iPSCs and ESCs were maintained in mTeSR1 medium (Stemcell Technologies) on Matrigel-coated plates (Corning, 354230) with daily medium replacement. We followed NIH 4D nucleome standard protocol for cell maintenance. HMC3 and SH-SY5Y were purchased from ATCC and were cultured in DMEM/F12 supplemented with 10% fetal bovine serum (FBS).

### Differentiation of H1-hESCs to NPCs and astrocytes

H1-hESCs derived NPCs and astrocytes were generated according to a previous report^79^. In brief, hESCs were dissociated into single cells and cultured in AggreWell 800 plates (Stemcell Technologies, 34811) to form embryoid bodies (EBs) in Neural Induction Medium (Stemcell Technologies). On day 5, EBs were collected and transferred to 6-well plates coated with Matrigel for culture in the NPC medium. Neural rosettes were collected and dissociated into single cells using Neural Rosette Selection Reagent (StemCell Technologies, 05832) after 14 days of culture. The hESCs-derived NPCs were then cultured in Neural Progenitor Medium (Stemcell Technologies, 05833), and passaged every 4-5 days. Following validation by immunostaining with NPC markers, the cells were cryopreserved in liquid nitrogen for future usage. For the differentiation of NPCs into astrocytes, NPCs were dissociated into single cells using accutase (Thermo Fisher, 00-4555-56) and plated on Matrigel-coated plates in astrocyte medium (ScienCell Research Laboratories, 1801) for 20 days, followed by splitting and subculturing until day 75. After validation by immunostaining with astrocyte markers, the cells were frozen for future analysis.

### Differentiation of WTC11 iPSCs to microglia

iPSC-derived microglia were generated following established protocols with some modifications^80,81^. In brief, iPSCs were cultured in mTeSR plus medium (Stemcell Technologies) on Matrigel-coated plates with daily medium replacement. The cells were dissociated into single cells using accutase and cultured in a 96-well ultra-low attachment plate (Corning, 7201680) to form yolk-sac embryoid bodies (YS-EBs) in 100 µL YS-EBs medium (10 mM ROCK inhibitor, 50 ng ml-1 BMP-4, 20 ng ml-1 SCF, and 50 ng ml-1 VEGF-121 in mTeSR plus), with 10,000 cells per well on Day 0. The culture medium was replaced on Days 2 and 4 with fresh YS-EBs medium without ROCK inhibitor. On Day 5, the YS-EBs were transferred to a 6-well plate with 10-12 YS-EBs per well and incubated in 3 mL hematopoietic medium (2 mM GlutaMax, 100 U ml-1 penicillin, 100 mg ml-1 streptomycin, 55 mM β-mercaptoethanol, 100 ng ml-1 M-CSF, and 25 ng ml-1 IL-3 in X-VIVO 15) to induce the generation of primitive macrophage precursors (PMPs). The medium (2 mL) was replaced with fresh hematopoietic medium every 5 days, and PMPs typically emerged as rounded brilliant cells in suspension within 2 weeks. Subsequently, PMPs were harvested from suspension and cultured on Matrigel-coated 6-well plates in microglia medium (DMEM/F12, 2X insulin-transferrin-selenium, 2X B27, 0.5X N2, 1X GlutaMax, 1X non-essential amino acids, 400 μM monothioglycerol, 5 μg ml-1 insulin) supplemented with three protein factors (100 ng ml-1 IL-34, 50 ng ml-1 TGFβ1, and 25 ng ml-1 M-CSF) for 25 days, with half-medium replacement every 3 days. After 25 days of culture, the cells were further incubated in microglia medium supplemented with five protein factors (100 ng ml-1 IL-34, 50 ng ml-1 TGFβ1, 25 ng ml-1 M-CSF, 100 ng ml-1 CD200, and 100 ng ml-1 CX3CL1) for an additional 3 days to achieve further maturation. Following validation through immunostaining with microglial markers, the iPSC-derived microglia were ready for subsequent treatment or analysis.

### i3Neuron differentiation

WTC11-Ngn2 iPSCs were utilized to generate i3Neurons following a previously established protocol ^52^. For pre-differentiation, WTC11-Ngn2 iPSCs were cultured with 2 μg ml-1 doxycycline in knockout DMEM/F12 medium supplemented with 1× N2, 1× NEAA, 1 μg ml−1 mouse laminin, 10 ng ml−1 brain-derived neurotrophic factor (BDNF) and 10 ng ml−1 NT3 for three days. ROCK inhibitor (10 μM) was included in the predifferentiation media for the first day, and the medium was replaced daily. During the maturation phase, predifferentiated neural cells were dissociated into single cells using accutase and plated on Poly-L-Ornithine-coated plates in maturation medium containing equal parts DMEM/F12 and Neurobasal-A with 2 μg ml−1 doxycycline, 0.5× B-27, 0.5× N-2, 1× NEAA, 0.5× GlutaMax, 1 μg ml−1 mouse laminin, 10 ng ml−1 BDNF and 10 ng ml−1 NT3. Doxycycline was only added to the maturation medium on the first day and omitted thereafter. Half of the medium was replaced weekly for the first 2 weeks, and the medium volume was doubled on week 3. Subsequently, one third of the medium was replaced weekly until 8 weeks for further treatment.

### *MAPT-paRNA* stability measurement

To study the effect of m6A methylation on *MAPT-paRNA* stability, 8-week-old i3Neurons were treated with m6A/METTL3 inhibitor STM2457 (10 µM, MedChemExpress, HY-134836) or DMSO vehicle for 24 hours. Following this treatment, Flavopiridol (2 µM, Sigma, F3055) was added to inhibit transcriptional activity. The RNA levels of *MAPT-paRNA* were then measured at various time points after transcriptional inhibition using RT-qPCR.

### Antisense oligonucleotides (ASOs)-based Knockdown

Knockdown experiments were performed using antisense oligonucleotides (ASOs) with modifications by the Affinity Plus nucleic acid modification (IDT) (**see Supplementary Table 9**). A sequence-scrambled ASO (Scr ASO) was used as a control. ASOs was dissolved in TE buffer (10 mM Tris-HCl, pH 7.4, 1 mM EDTA) and directly added to the i3Neuron culture without any transfection reagent (termed gymnosis) at a final concentration of 10µM. After 3 days of ASOs treatment (unless otherwise indicated), cells were harvested for downstream analysis.

### Western blotting analysis

Cells were washed twice with cold PBS buffer and subsequently lysed in RIPA buffer (50 mM Tris, pH 7.4, 150 mM NaCl, 1 mM EDTA, 0.1% SDS, 1% NP-40, 0.5% sodium deoxycholate) containing cOmplete Mini Protease Inhibitor Cocktail (Roche, 11836153001) on ice for 30 minutes. Following centrifugation at 12,000 × g for 10 minutes at 4°C, the supernatants were combined with 2x Laemmli sample buffer (Bio-Rad, 1610737) and boiled at 95 °C for 10 minutes. The protein samples were loaded and separated using 4%–15% SDS–PAGE gradient gels, then transferred to a PVDF membrane (Bio-Rad, 1620260). Subsequently, the membranes were blocked in 5% skim milk in TBST (20 mM Tris, 150 mM NaCl, and 0.2% Tween-20, w/v) for 1 hour and incubated with primary antibodies, anti-SYNGAP1 (1:2000; CST; #62193S), anti-MEF2C (1:1000; Abcam; #ab211493), anti-HDAC4 (1:2000; CST; #7628T), anti-Synapsin1 (1:1000; SYSY; #106011), anti-PSD95 (1:1000; Sigma; #MAB1596), anti-GluN1 (1:1000; SYSY; #114011), anti-GluA2 (1:1000; BioLegend; #810501), anti-Total Tau (1:20000; Dako; #A0024), anti-pTau (Ser202/Thr205) (1:1000; Thermo Fisher; #MN1020), anti-Tubulin (1:5000; Thermo Fisher; #A11126), or anti-GAPDH (1:50000; Proteintech; #60004). After three washes in TBST, the membranes were incubated with horse-radish peroxidase (HRP)-conjugated secondary antibody for 1 hour. Following six washes, protein bands were visualized using the Enhanced Chemiluminescence kit (Pierce) and developed in the Bio-Rad ChemiDoc gel imaging system. Band intensities were quantified using ImageJ software, with tubulin blots serving as loading controls.

### Glutamate-induced neuronal excitotoxicity assay

To investigate the neuroprotective role of *MAPT-paRNA*, we implemented glutamate-induced excitotoxicity based on established protocols with slight modifications^74,82^. In detail, i3Neurons were cultured and differentiated in a 24-well plate for 8 weeks and treated with Scr ASO or MAPT-paRNA-ASO1 for 3 days. Subsequently, vehicle or 10µM L-glutamate (Sigma, G1251) was introduced into the medium for 2 hours. Then the cells were stained with Hoechst 33342 (5 μg ml−1) (Thermo Fisher, 62249) and propidium iodide (PI, 1 μg ml−1) (Thermo Fisher, P21493) for 30 min to visualize total and dead cells, respectively. Images were captured using a Keyence BZ-X810 Fluorescence Microscope, and neurons from at least 3 wells were quantified. To assess the toxicity of ASOs alone on i3Neurons, the same protocol was applied, excluding the L-glutamate treatment.

### Beta-amyloid (Aβ-42)-induced cytotoxicity

To assess the role of MAPT-paRNA under Aβ-induced stress, 8-week-cultured i3Neurons were treated with either Scr ASO or MAPT-paRNA-ASO1 in the presence of scrambled beta-amyloid (Scr Aβ-42, Anaspec, #AS-25382) or beta-amyloid (Aβ-42, Anaspec, #AS-64129-05) for six consecutive days. Cell cytotoxicity was evaluated daily by measuring lactate dehydrogenase (LDH) levels in the culture media using the LDH Assay Kit (Sigma, #MAK066).

### Electrophysiological recordings

Electrophysiological recordings were conducted at room temperature using 8-week-cultured i3Neurons in a whole-cell patch-clamp configuration. Patch pipettes with a resistance of 8–15 MΩ were filled with an internal solution containing 135 mM CsF, 33 mM CsCl, 2 mM MgCl₂, 1 mM CaCl₂, 11 mM EGTA, and 10 mM HEPES (pH 7.4). The external ACSF solution consisted of 140 mM NaCl, 7 mM KCl, 2 mM CaCl₂, 10 mM HEPES, 10 mM glucose, 10 μM bicuculline, and 1 mM TTX (pH 7.4). To selectively study NMDA receptor-mediated currents with glutamate, non-NMDA receptors were blocked using 10 μM NBQX. Conversely, for investigating AMPA receptor-mediated currents, NMDA receptors were blocked with 100 μM APV and 25 μM DCKA. External solutions were locally applied to neurons using an SF-77B Fast-Step perfusion system (Warner Instruments). All recordings were performed at room temperature with a holding potential of −60 mV using an Axopatch 200B amplifier (Molecular Devices). Data were acquired at 10 kHz using pCLAMP10.7 software (Molecular Devices) and filtered online at 5 kHz. Current densities were calculated by dividing the recorded current (pA) by the measured cell membrane capacitance (pF).

### Immunocytochemistry

For immunofluorescence immunostaining, the cells were cultured on coverslips, then washed twice with PBS and fixed in cold methanol (−20°C) for 15 min. Subsequently, the cells were permeabilized and blocked with PBS containing 5% goat serum, 5% BSA (bovine serum albumin), and 0.3% Triton X-100 for 1 h, followed by incubation with primary antibodies diluted in blocking buffer overnight at 4°C. After incubation with fluorescence-conjugated secondary antibodies at room temperature for 1 hour, cell nuclei were counterstained with DAPI, and images were acquired using a confocal microscope (Nikon A1) or fluorescence microscope (Keyence). Antibodies used for immunocytochemistry were against PSD95 (1:200; Sigma; #MAB1596), GluN1 (1:200; SYSY; #114011), NeuN (1:200; ThermoFisher; #702022), and MAP2 (1:400; Abcam; #ab5392). To quantify the intensity of immunostaining images, ImageJ software (ImageJ website: http://imagej.nih.gov.laneproxy.stanford.edu/ij/) was used. A minimum of 15 neurons from 3 separate experiments were quantified for statistical analysis.

### Quantitative RT-PCR (qRT-PCR)

Total cellular RNAs were extracted by TRIzol (Thermo Fisher) or Zymo RNA miniprep kit (Zymo Research), following the respective manufacturer’s instructions. The extracted RNA underwent reverse transcription by SuperScript IV (Thermo Fisher) for first strand cDNA synthesis using random hexamer. For qPCR, the SsoAdvanced Universal SYBR green Supermix (Bio-Rad) was used according to the standard parameters recommended by the manufacturer. Primer sequences were shown in **Supplementary Table 9**. All qPCR analyses were performed in triplicate, and each value was denoted by a black dot.

### RNA-seq, MeRIP-seq, and MeRIP-qPCR

RNA-seq and MeRIP-seq were performed according to our previous report with some modifications ^12^. Total RNA (2-5 μg) was fragmented at 70 °C for 6 min in the fragmentation buffer (10 mM ZnCl2, 10 mM TrisHCl pH7.4), following precipitation the RNA was resuspended with ice-cold IP buffer (10 mM Tris-HCl, pH 7.4, 150 mM NaCl, 0.1% NP-40, 20 U/mL Superase-In, 1× Protease Inhibitor) to a final volume of 1050 μL. Of this, 50 μL of diluted RNA was utilized for RNA-seq (5% input for m6A IP) and the remaining 1000 μL was reserved for MeRIP-seq (Methylated RNA immunoprecipitation sequencing). To prepare anti-m6A antibody conjugated beads, 20 μL of Dynabeads Protein G beads (Thermo Fisher) was washed 3 times with 1ml IP buffer and then incubated with 1 μg of m6A antibody (Synaptic Systems, # 202003) at room temperature for 30min. The conjugated anti-m6A-Dynabeads were washed in 1ml IP buffer 3 times. Subsequently, the RNA sample was mixed with the beads and rotated at 4 °C for 3 hours. The beads were then washed in the IP buffer 5 times. The immunoprecipitated RNAs were extracted from the beads by TRIzol-LS (Thermo Fisher). The extracted RNA was subjected to the next generation sequencing library preparation for RNA-seq and MeRIP-seq, as well as qPCR, to quantify the m6A levels.

### Library Preparation and Sequencing

RNA samples were collected by one of two methods: lysis in TRIzol reagent (Thermo Fisher) for RNA-seq and MeRIP-seq, followed by standard phenol-chloroform extraction methods; or alternatively, lysis was performed using Zymo Research Quick-RNA MiniPrep kit (Genesee Scientific) for regular qPCR following the manufacturer protocol. Libraries for RNA-Seq and MeRIP-Seq were made with NEBNext Ultra II Directional Library Prep Kit for Illumina (NEB, #E7760) following the manufacturer’s instructions. Ribosome RNA was depleted with NEBNext rRNA Depletion Kit (NEB, #E6301S). The generated libraries were quantified using the Qubit-3 system and/or qPCR. Sequencing of the libraries was performed using a NextSeq 550 platform (Illumina) with paired-end sequencing (40/40nt mode).

### RNA-seq data analysis

We used FastQC (v.0.11.8) (https://www.bioinformatics.babraham.ac.uk/projects/fastqc/) to check the quality of RNA-seq raw reads. RNA-seq clean reads were aligned to the human reference genome (hg19) or mouse reference genome (m31) from the UCSC database using HISAT2 (v.2.2.1)^83^ with *-k 1 --no-discordant --rna-strandness RF*. StringTie (v.2.1.6)^84^ was implemented to assemble and quantify transcripts based on gencode.v19.annotation.gtf (human) or gencode.vM31.annotation.gtf (mouse). DESeq2 (v.1.34.0)^85^ was run to analyze differentially expressed genes (DEGs). DEGs between AD and Normal groups were defined based on *p.adj* < 0.05 and absolute log_2_foldchange > 1. Differentially expressed paRNAs between Normal and AD/ALS/PD/FTD brains were selected based on *p.adj* < 0.05 and absolute log_2_foldchange > log2(1.5). For *MAPT-paRNA* control and KD RNA-seq data, DEGs were defined based on *p.adj* < 0.05 and absolute log_2_foldchange > log2(1.5).

### ChIP-seq analysis

We firstly downloaded Non Cognitive Impairment (NCI) and AD H3K27ac and H3K4me3 ChIP-seq fastq files from ENCODE (https://www.encodeproject.org/brain-matrix/?type=Experiment&status=released&internal_tags=RushAD) and H3K27ac ChIP-seq data (GSE153875) (Nativio et al. ^36^). We used FastQC (v.0.11.8) (https://www.bioinformatics.babraham.ac.uk/projects/fastqc/) to check the quality of ChIP-seq reads. We applied Bowtie2 (v.2.2.5)^86^ with --very-sensitive to map clean reads from ChIP-seq to hg19 from the UCSC database, and duplicates are further removed with picard. MACS3 (v.3.0.0a6)^39^ (https://github.com/macs3-project/MACS) was used to call H3K27ac and H3K4me3 peaks, respectively.

### De novo transcript calling

We followed the previous method^42,43^ to identify RNA transcripts using STAN R package (v.2.22.0)^87^. Briefly, we merged the bam files from the same group to increase the sequencing depth, and convert the bam to bed files. Next, we applied the STAN package to the bed file with default parameters. To define paRNAs, the transcription start sites (TSSs) of any possible paRNAs overlapping with the promoter regions (−2kb to 2kb) of genes were retained.

### Long-reads Nanopore Direct RNA-seq data Analysis

We downloaded the long-reads RNA-seq datasets from ENCODE (https://www.encodeproject.org/brain-matrix/?type=Experiment&status=released&internal_tags=RushAD). We followed the ENCODE pipeline to reanalyze long-read RNA-seq datasets. Here we used minimap2 (v.2.17-r941)^88^ to map the long-read RNA-seq data to the hg19 genome build. Next, TALON (v5.0)^89^ was used to quantify the expression of transcripts. Any lincRNA, antisense RNA, processed_transcript or unannotated transcripts overlapping with the transcription start sites (TSSs) of PCGs were retained.

### MeRIP-seq data analysis

Similar to RNA-seq data analysis, we used the same pipeline to map clean MeRIP-seq data to the human reference genome. We used MACS3 (v.3.0.0a6)^39^ (https://github.com/macs3-project/MACS) with --nomodel --extsize 100 to call m6A peaks.The high-confidence m6A peaks were retained based on q value ≤ 10^−5^ and fold change ≥ 5. We used bedtools (v.2.30.0)^90^ to get intersected m6A peaks detected in at least 3 samples. We used ChIPseeker (v.1.40.0)^91^ to annotate the genomic region of a m6A peak as TSS, exon, 5′ UTR, 3′ UTR, intronic or intergenic. The ‘gencode.v19.annotation.gtf’ was used as the transcript database. Gene promoter regions were defined on the basis of -2 kb distance from TSS. For m6A peaks annotated to ncRNAs, we used m6A peaks to overlap with ncRNAs identified by *de novo* calling or repetitive elements downloaded from UCSC Genome Browser.

### Differential analysis of m6A peaks

The gained or lost m6A peaks were determined by whether the genomic coordinates of any m6A peaks overlap or not. Of note, this criterion is markedly biased by the peak calling algorithm. For example, the enrichment fold of a genomic region is 4.9 and 5.1 in Normal and AD brains, respectively. Thus, this genomic region would have a m6A peak in AD, but not in Normal brains. Moreover, the difference in m6A ratio (MeRIP/input) in this genomic region between Normal and AD brains was not large. The existing algorithms analyzing differential m6A peaks divided the genomic regions into different bin sizes and compared the differences in m6A ratio between case and control samples. These algorithms have at least 3 limitations. First of all, there is no gold standard for choosing a proper bin size. In addition, continuous bin sizes have significant differences in m6A ratio, these continuous bin sizes would be concatenated, which would make a m6A peak region large. Third, the genomic coordinates of these differential m6A peaks called by the existing algorithms do not match the genomic coordinates of m6A peaks called by MACS3. To address these questions, we merged the m6A peaks from Normal and AD brains to obtain unique m6A peaks, and then calculated the RNA (referred to as *raw FPKM(RNA)*) and m6A methylation (referred to as *raw FPKM(m6A)*) levels for each m6A peak region. Given the sparsity of RNAs in some m6A peak regions, we added a pseudocount of the first quantile of RNA levels from each input sample to *raw FPKM(RNA)* to get the *final FPKM(RNA)*, and calculated the m6A ratio based on the *raw FPKM(m6A)* divided by *the final FPKM(RNA)*. Here is the formula for the normalized m6A ratio: 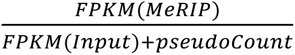. We further removed the outliers of m6A ratio less than 3.5 based on the quantile distributions of m6A ratio. Next, we performed the paired student’s t test for m6A ratio of each m6A peak between AD and Normal brains. We selected the differential m6A peaks based on the fold change > 1.5 and p < 0.1.

### Gene Ontology analysis

We applied gProfiler2 (v.0.2.2)^92^ to analyze the functional enrichment of genes. Significant GO terms will be selected based on a 5% false discovery rate (FDR).

### snRNA-seq data analysis

We downloaded control and Alzheimer’s disease snRNA-seq datasets^52^. Based on the previous method^93^, we used the cellranger count (v.7.0.0)^94^ with default parameters to align the snRNA-seq data to the hg19 human reference genome including mRNAs and paRNAs to produce the raw cell-by-gene count matrix based on the barcode matrix for all snRNA-seq libraries. Next, we used Seurat (v.5.0.0)^95^ to filter out low quality nuclei based on nFeature_RNA > 200 & nFeature_RNA < 10000 & percent.mt < 5. Subsequently, the merged expression matrix was normalized by the NormalizeData with default parameters from Seurat. The principal components were calculated using the first 3000 variable genes, and the Uniform Manifold Approximation and Projection (UMAP) analysis was performed with RunUMAP from Seurat. The FindAllMarkers from Seurat was used to analyze the differentially expressed genes across different clusters. Finally, we used the previously identified marker genes^52^ to define the different cell clusters or cell types. Here if a paRNA was detected in more than 3% cells of a given cell type, then that paRNA was defined as a detected paRNA in that cell type in snRNA-seq data.

### MUSIC data analysis

To analyze how many paRNAs are detected by MUSIC data, we directly used the processed reads/cells/complex matrix reported in Wen et al.^59^, (which is in hg38 genome build). We liftovered the genomic coordinates of our paRNAs from hg19 genome to hg38, and intersected the genomic coordinates of paRNAs we identified in human brain bulk RNA-seq datasets with the RNA reads from MUSIC data. Here if a paRNA was detected in more than 3% cells of a given cell type based on the cell barcoder (CB), then that paRNA was defined as a detected paRNA in that cell type in MUSIC data. The expression pattern of paRNAs is similar when using a more stringent cutoff (e.g., 5% of cells expressing paRNAs). To identify the target genes of paRNAs of interest, the DNA reads interacting with paRNAs were identified based on shared cell barcodes and molecular barcodes (CBMB) between RNA reads and DNA reads. To annotate these DNA reads to genes, we used the gene body to overlap with the DNA reads interacting with paRNAs. To remove technical issues, we remove large molecular complexes with RNA reads>50 and DNA reads>1000. We followed the method described by Wen et al.^59^ to calculate the contact strength between RNA reads and DNA reads in a complex: 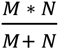, M and N represent the number of RNA reads and DNA reads in a complex, respectively. Synaptic genes (n=393) were extracted from **Supplementary Fig, 11C**, and then we calculated the contact strength between AD-deregulated paRNAs (DE paRNAs, n = 47) / non-AD-deregulated paRNAs (n = 346) and these synaptic genes (n=393).

### Motif analysis

We ran Homer (findMotifsGenome -rna -len 5,6,7)^96^ to analyze m6A peaks and to identify m6A motifs.

### Quantification and statistical analysis

All bar graphs of qPCR data were presented as mean ± SD as indicated in figure legend using Prism 6.0. Two-tailed Student’s t-test was used to compare means between groups as indicated; p < 0.05 was considered significant, and p values with asterisks are indicated in each figure panel. ∗, p < 0.05; ∗∗, p < 0.01; ∗∗∗p < 0.001. Statistical details of experiments can be found in the figure legends. No statistical methods were used to pre-determine sample sizes. For all boxplots, the central lines represent medians; box limits indicate the 25th and 75th percentiles; and whiskers extend 1.5 times the interquartile range (IQR) from the 25th and 75th percentiles. For NGS data analysis, statistical analysis was performed using Python (v.3.8) and R (v.4.3), and is indicated in the legend of each figure panel.

## Acknowledgments

This work was supported the University of Texas UTHealth McGovern Medical School at Houston Startup fund, NIH/NIA (R01AG082132), the ‘4D Nucleome’ program (U01HL156059), NIGMS (R01GM136922), Cancer Prevention and Research Institute of Texas (CPRIT, RR160083), and Welch foundation (AU-2000-20220331) to W.L.; by NIH grant U01DA052769 to X.X.; by NIH/NCI (R01CA246130), NIH/NHLBI (R01HL142704), and CPRIT(RR160019) to D.-F.L.; by grants from ‘4D Nucleome’ program (U01CA200147), NIGMS (R01GM138852), NIDDK (DP1DK126138), NICHD (R01HD107206), and a Kruger research grant to S.Z.; by grants from NIH RF1 AG020670, RF1 NS093652, and P01 AG066606 to Z.H.; by NIGMS R35GM122528 to V.J., and by NIAID P01AI077774-S and NIA RF1AG059321 to C.S.. B.H. was partially supported by the CPRIT Biomedical Informatics, Genomics and Translational Cancer Research Training Program (BIG-TCR) postdoctoral fellowship (RP210045). Our next generation sequencing work was mostly conducted with the UTHealth Cancer Genomics Core, which received funding from CPRIT (RP240610). We thank WiCell and Coriell for hESCs and iPSCs, and thank Drs. Li Gan (Weill Cornell) and Yin Shen (UCSF) for generously sharing the WTC11-ngn2 cell model. We are grateful to the donors who contributed samples and knowledge to the ENCODE, RUSH program and AD Knowledge Portal and for kindly sharing these datasets with the scientific community.

## Author contributions

W.L., B.H., and Y.S. conceived the project. B.H. conducted all the dry lab experiments. Y.S. did most of the wet lab experiments, with help from Y.-T.C., X.Z. and N.D. F.X. generated the original RNA-Seq and MeRIP-Seq datasets. H.Z. and R.C. acquired human brain tissues and contributed to i3Neuron work, and D.-F.L. and C.S. helped on neuronal/glial cell differentiation and culture. X.W. and S.Z. shared and helped with MUSIC data. E.F. and V.J. contributed the electrophysiology results. W.L., B.H., and Y.S. wrote the manuscript, with inputs from all authors. These authors contributed equally: Benxia Hu, Yuqiang Shi.

## Data availability

Public data:

H3K4me3 and H3K27ac ChIP-seq, and RNA-seq datasets generated from No Cognitive Impairment and AD DLPFC are available through (https://www.encodeproject.org/brain-matrix/?type=Experiment&status=released&internal_tags=RushAD). Old and AD lateral temporal lobe RNA-seq datasets are available through GSE104704 (Nativio et al., PMID: 32989324), and H3K27ac ChIP-seq data are available through GSE153875 (Nativio et al., PMID: 29507413). RNA-seq data generated from the 5xFAD mice brain (right cerebral hemisphere) are available through https://www.synapse.org/#!Synapse:syn21983020. RNA-seq data generated from the 3xTg-AD mouse mode are available through https://www.synapse.org/#!Synapse:syn22964719. RNA-seq data generated from the P301S mouse model are available through GSE226385 (Udeochu et al., PMID: 37095396). RNA-seq data generated from Parkinson’s disease are available through GSE148434 (PD, Lee et al., PMID: 37058563), Frontotemporal Dementia are available through GSE116622 (FTD, Conlon et al., PMID: 30003873), and Amyotrophic lateral sclerosis are available through GSE124439 (ALS, Tam et al., PMID: 31665631). RNA-seq data from iPSC-derived cellular models are available through GSE102956 (Lin et al., Neuron, PMID:29861287). DLPFC Normal and AD snRNA-seq data are available through GSE174367 (Morabito et al., PMID:34239132).

Data generated by this work:

Data generated in this study have been deposited to NCBI GEO (GSE266459). The reviewer token is *<u>mjidasquplmzbcx</u>*. Source Data are provided with this paper (Source Data Files 1,2,3). Source data and codes in this paper are also available at Zenodo (https://doi.org/10.5281/zenodo.15019230)^97^.

## Code availability

All custom code used in this work is available in the following github repository:(https://github.com/BenxiaHu/MAPT-paRNA_in_AD). Genomic tracks are generated by the HiCPlot NGStrack function (https://github.com/BenxiaHu/HiCPlot).

