## Supplementary information for "Rewired m6A methylation of promoter antisense RNAs in Alzheimer’s disease regulates global gene transcription in the 3D nucleome"

### Supplementary Figures and Legends

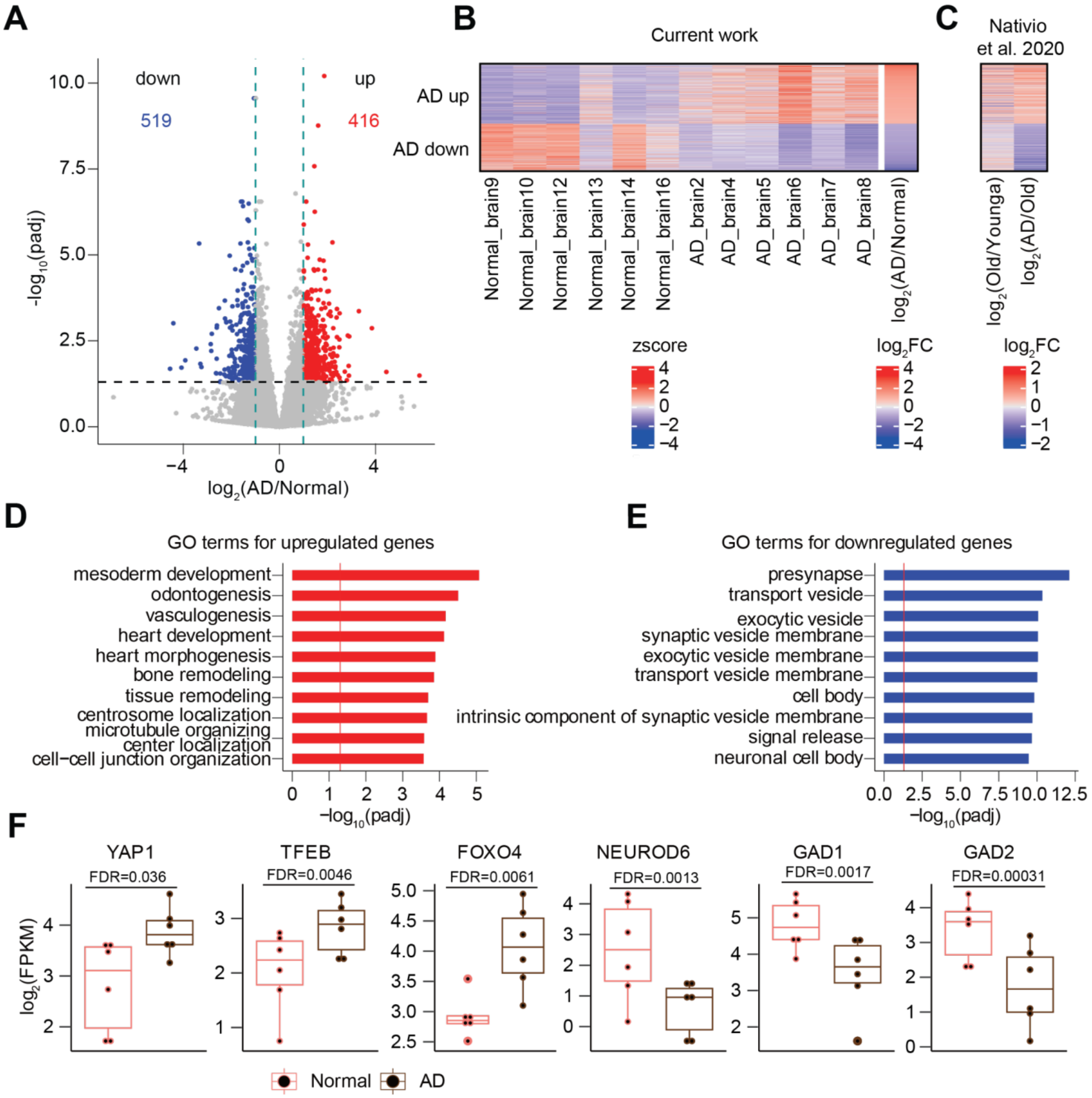


**Supplementary Figure 1. The alterations of transcriptome in the human AD brains. A**. A volcano plot showing differentially expressed genes between Normal and AD brains. Red and blue dots represent upregulated and downregulated genes in AD, respectively. **B**. A heatmap showing z-score-transformed expression levels of differentially expressed genes between Normal and AD brains from our samples, and from Nativio et al.[36]. **C**. A heatmap showing that the deregulated genes in AD (as in B panel) are not seen to be deregulated in normal aging, revealed by re-analysis of RNA-seq data from Nativio et al.[36]. **D-E**. GO analyses for upregulated and downregulated genes in AD, respectively. The red lines represent padj values at 0.05. **F**. Boxplots showing the expression levels of several examples of differentially expressed genes, some of which play important roles in neurodevelopment, AD disease or synapse biology. FDR values were calculated by DESeq2 and shown on top. Boxplots indicate the interquartile range with the central line representing the median, and the vertical lines extending to the extreme values in the group.

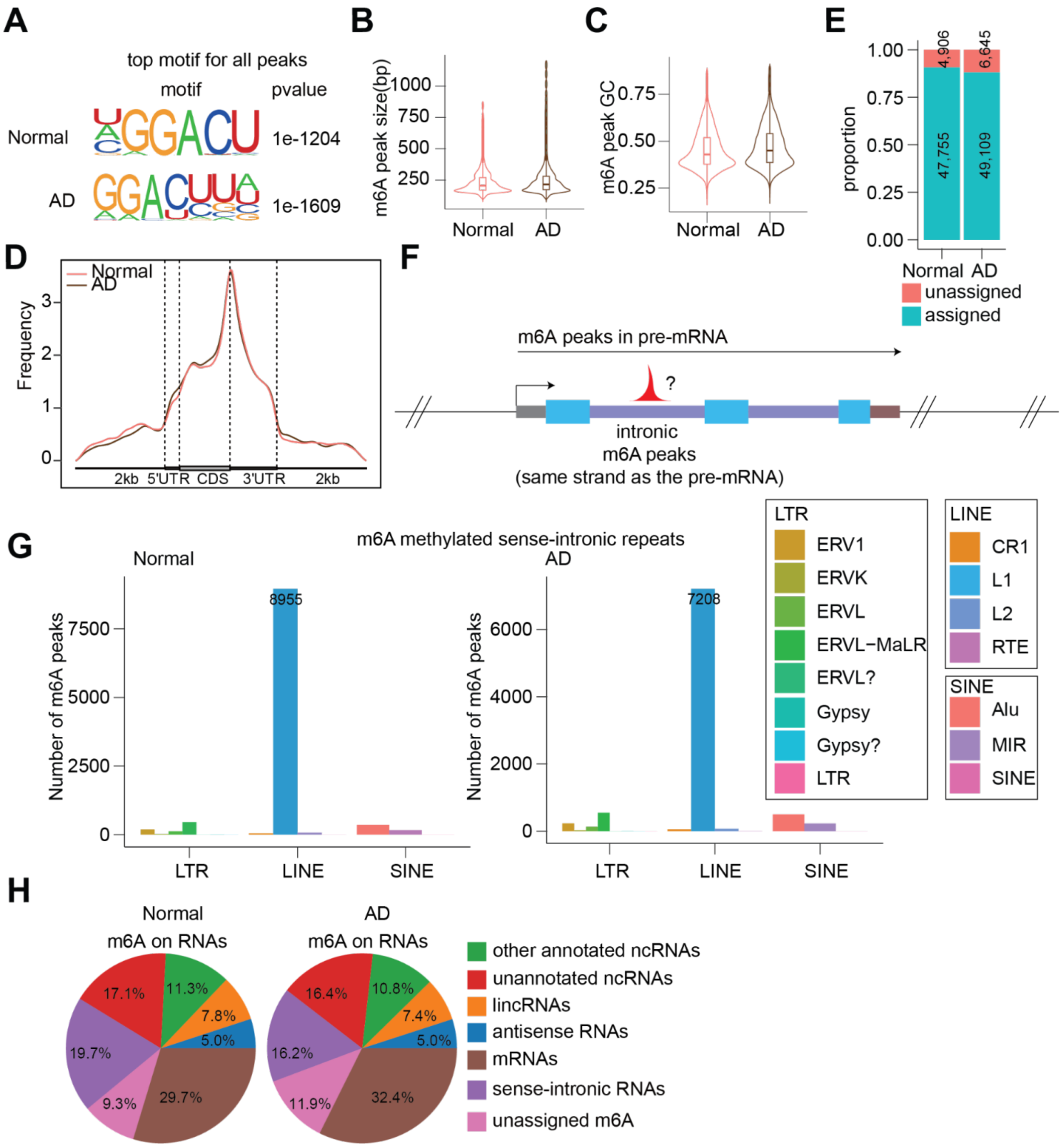


**Supplementary Figure 2. The characteristics and locations of m6A peaks in the human AD brains. A**. Motif analysis of all m6A peaks detected in the Normal and AD brains. **B-C**. Violin plots showing the peak sizes and GC content of m6A peaks detected in the Normal and AD brains, respectively. **D**. Metagene profiles of m6A peak density along protein-coding gene transcripts with three non-overlapping segments (5' UTR, CDS, and 3' UTR) for the Normal and AD brains. **E**. The numbers of m6A peaks that are or are not assigned to any de novo transcripts in Normal and AD conditions, respectively. **F**. A diagram depicting examples of m6A peaks in gene introns that are included in the category of pre-mRNA peaks. **G**. The numbers of m6A peaks overlapping retrotransposon elements in intronic regions of pre-mRNAs in the Normal and AD brains, respectively. The categories of LTR, LINE and SINE were further divided into sub-groups. **H**. The distribution of m6A peaks annotated to various de novo called transcripts in the Normal and AD brains, respectively. Distinct from **Fig. 1D**, here pre-mRNA is broken down to two groups: mRNAs (5' and 3' UTRs and exons) and sense-intronic RNAs. The sense-intronic RNAs were not called as independent transcripts by de novo calling, but they are considered noncoding regions in our m6A peak definition as they may play regulatory roles in affecting gene transcription.

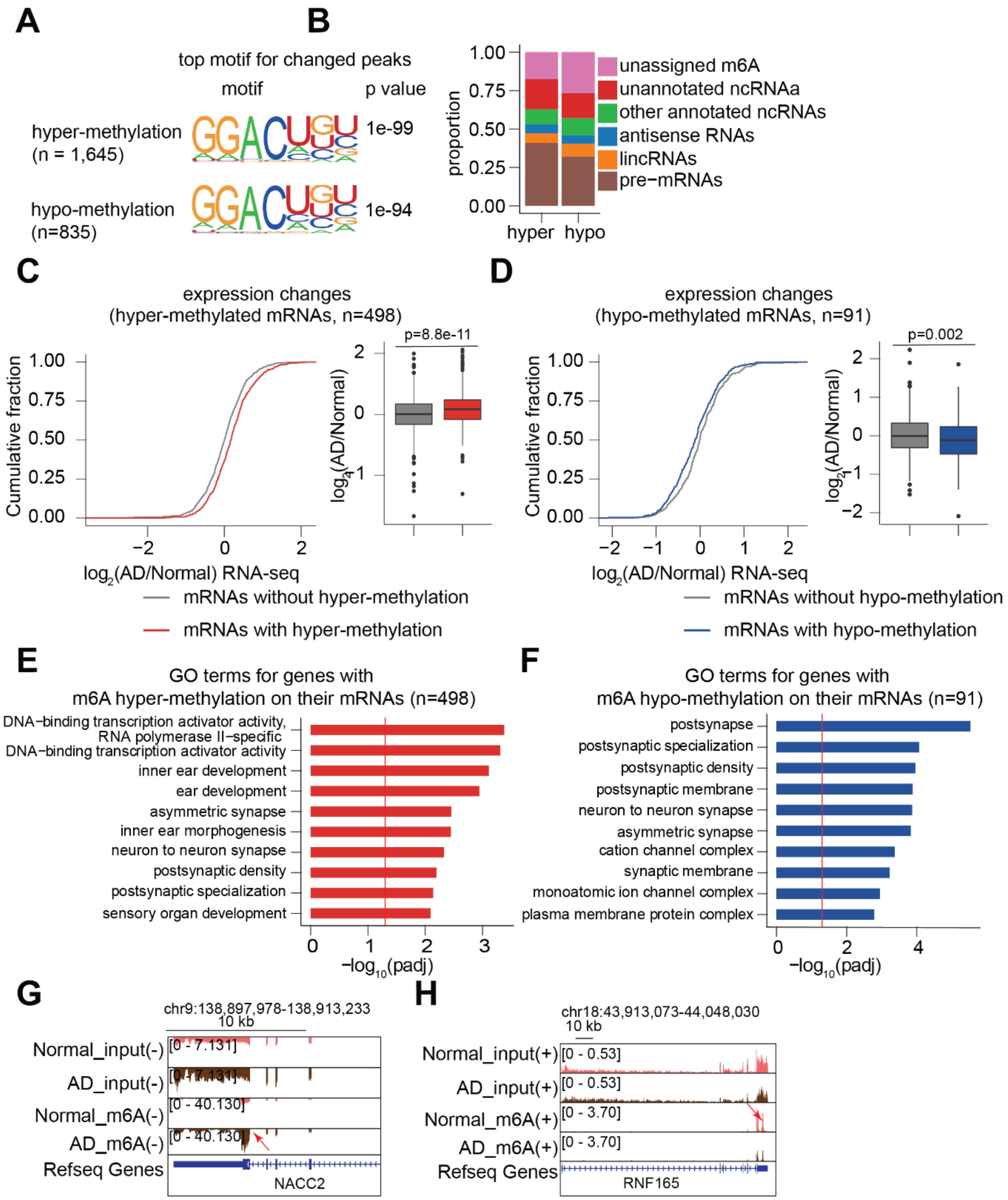


**Supplementary Figure 3. Landscape and alteration of m6A RNA methylome in mFC regions of human AD brains. A**. Motif analysis for hyper- and hypo-methylated m6A peaks detected in the AD brains. **B.** The distribution of AD-associated hyper- and hypo-methylated m6A sites to various ncRNAs. **C-D.** Cumulative distribution and boxplots of mRNA expression changes with or without hyper- and hypo-m6A-methylation. P values were calculated by a two-tailed non-parametric Wilcoxon–Mann–Whitney test. Box plots in the panels indicate the interquartile range with the central line representing the median, and the vertical lines extending to the extreme values in the group. **E-F**. GO analysis for genes whose mRNAs are associated with hyper- or hypo-methylated m6A peaks, respectively. The red lines represent padj values at 0.05. **G-H.** Genomic tracks show examples of differential m6A peaks between Normal and AD brains. Input and m6A represent RNA-seq and MeRIP-seq, respectively. (+) and (-) show the Watson and Crick strands, respectively. The red arrows indicate differential m6A peaks.

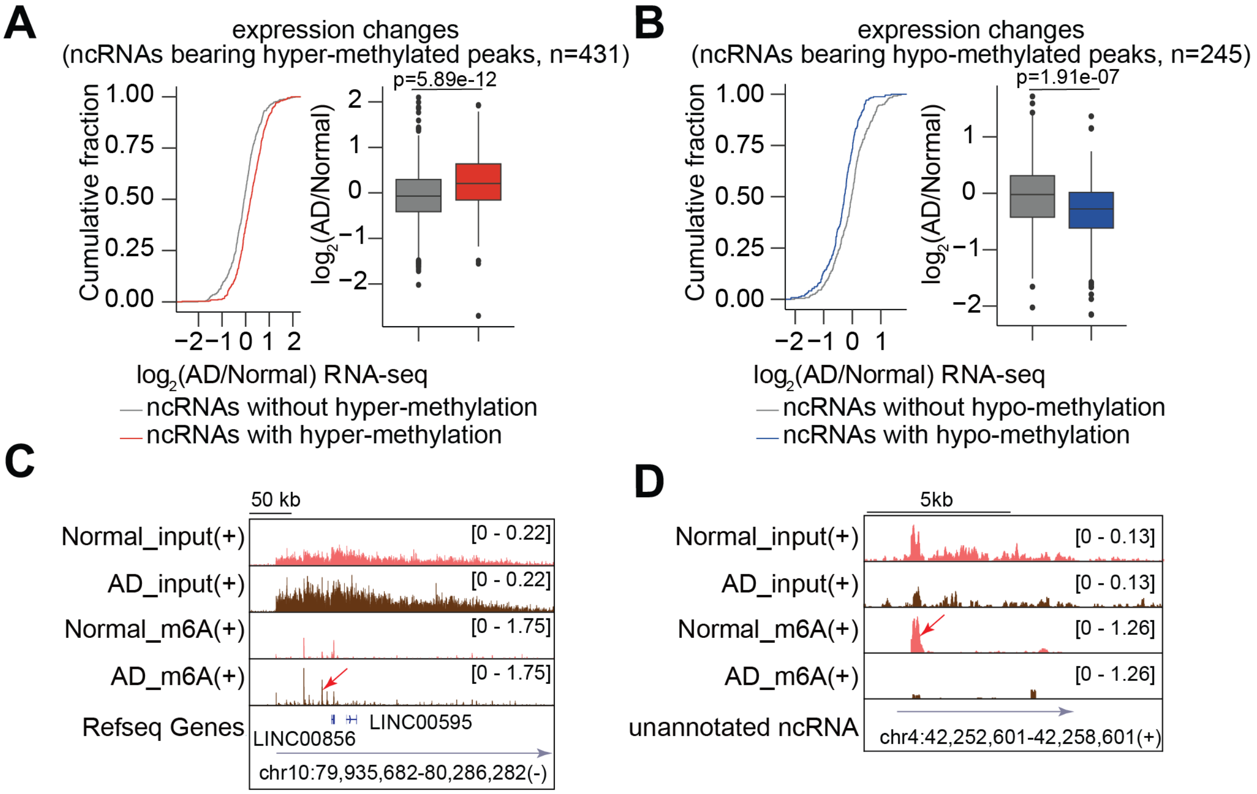


**Supplementary Figure 4. The correlation between differential m6A peaks and ncRNA expression in the human AD brains. A-B.** Cumulative distribution and boxplots of ncRNA expression changes with or without hyper- and hypo-m6A-methylation. P values were calculated by a two-tailed non-parametric Wilcoxon–Mann–Whitney test. Box plots in the panels indicate the interquartile range with the central line representing the median, and the vertical lines extending to the extreme values in the group. **C-D.** Genomic tracks showing examples of differential m6A peaks between Normal and AD brains. Input and m6A represent RNA-seq and MeRIP-seq, respectively. (+) indicates the Watson strand. The red arrow indicates differential m6A peaks.

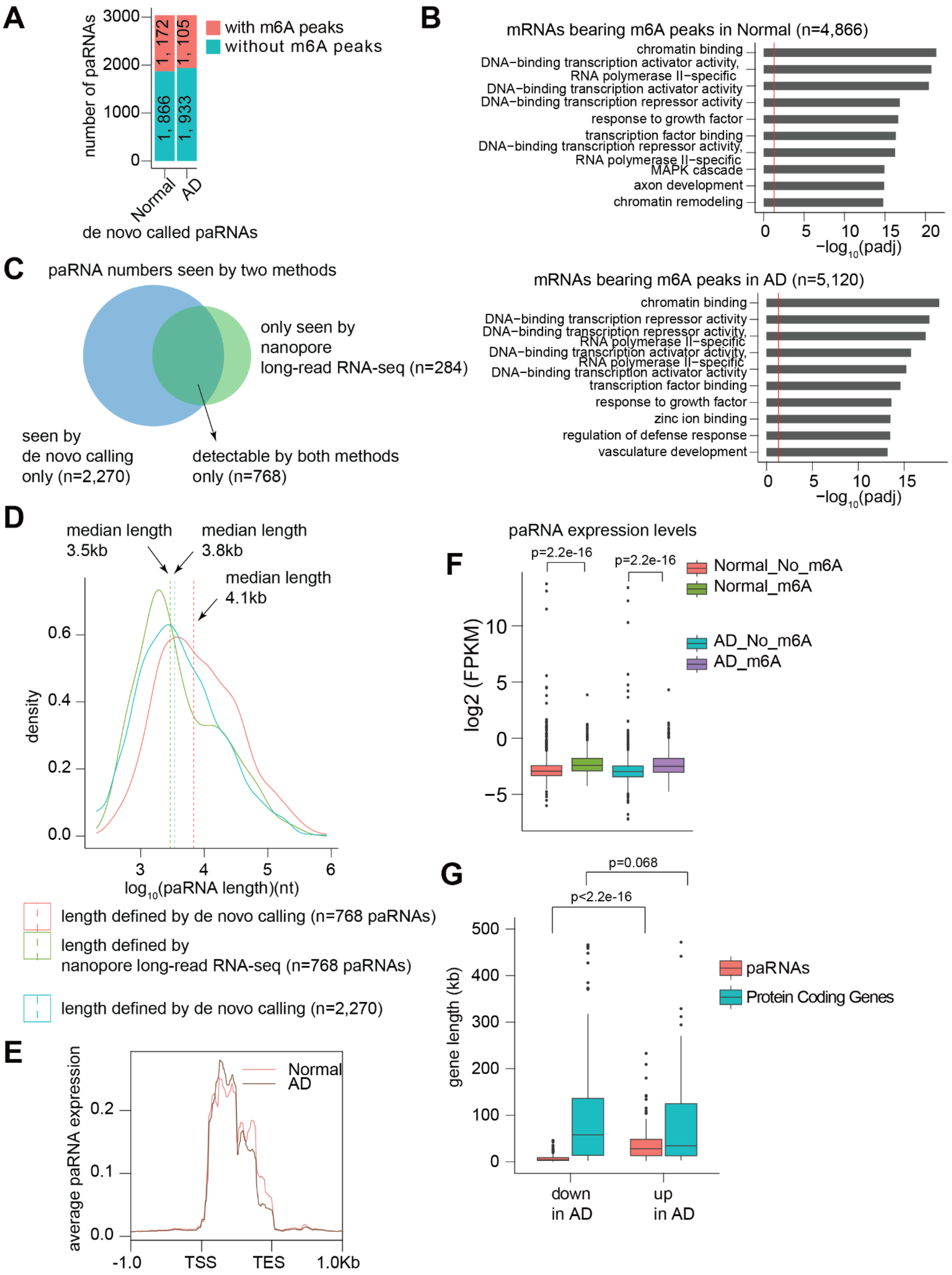
A
number of paRNAs
3000
2000
1000
0
1, 172
1, 105
1, 866
1, 933
Normal
AD
de novo called paRNAs
with m6A peaks
without m6A peaks
B
mRNAs bearing m6A peaks in Normal (n=4,866)
chromatin binding
DNA−binding transcription activator activity, RNA polymerase II−specific
DNA−binding transcription activator activity
DNA−binding transcription repressor activity
response to growth factor
transcription factor binding
DNA−binding transcription repressor activity, RNA polymerase II−specific
MAPK cascade
axon development
chromatin remodeling
−log10(padj)
mRNAs bearing m6A peaks in AD (n=5,120)
chromatin binding
DNA−binding transcription repressor activity
DNA−binding transcription repressor activity, RNA polymerase II−specific
DNA−binding transcription activator activity, RNA polymerase II−specific
DNA−binding transcription activator activity
transcription factor binding
response to growth factor
zinc ion binding
regulation of defense response
vasculature development
−log10(padj)
C
paRNA numbers seen by two methods
only seen by nanopore long-read RNA-seq (n=284)
seen by de novo calling only (n=2,270)
detectable by both methods only (n=768)
D
median length 3.5kb
median length 3.8kb
median length 4.1kb
density
log10(paRNA length)(nt)
length defined by de novo calling (n=768 paRNAs)
length defined by nanopore long-read RNA-seq (n=768 paRNAs)
length defined by de novo calling (n=2,270)
E
average paRNA expression
Normal
AD
-1.0
TSS
TES
1.0Kb
F
paRNA expression levels
p=2.2e-16
p=2.2e-16
log2 (FPKM)
Normal_No_m6A
Normal_m6A
AD_No_m6A
AD_m6A
G
p=0.068
p<2.2e-16
gene length (kb)
paRNAs
Protein Coding Genes
down in AD
up in AD

**Supplementary Figure 5. Validation of paRNAs using nanopore long reads sequencing, and their additional features**. **A**. A barplot showing the numbers of m6A peaks in the paRNAs. The red and blue bars represent m6A peaks overlapping and not overlapping with paRNAs, respectively. **B**. GO analyses for mRNAs having m6A peaks called in Normal and AD brains, respectively. The red lines represent padj value at 0.05. This serves as comparison to the GO patterns for paRNA-associated genes in **Fig. 3C,D**. **C**. A pie chart showing the numbers of paRNAs seen by the de novo calling with total RNA-seq data versus by nanopore long reads sequencing data. **D**. A density plot showing the length of paRNAs detected by the two different methods. **E**. A metaplot showing the average expression signals of paRNAs between Normal and AD brains (from our RNA-seq). **F**. Boxplots showing expression levels of paRNAs having or not having m6A peaks, in Normal or AD mFC regions, respectively. **G.** Boxplots showing the length of dysregulated paRNAs and protein-coding genes in AD brains. Red and blue boxplots represent paRNAs and protein-coding genes, respectively. P values were calculated by a two-tailed non-parametric Wilcoxon–Mann–Whitney test. Boxplots indicate the interquartile range with the central line representing the median, and the vertical lines extending to the extreme values in the group.

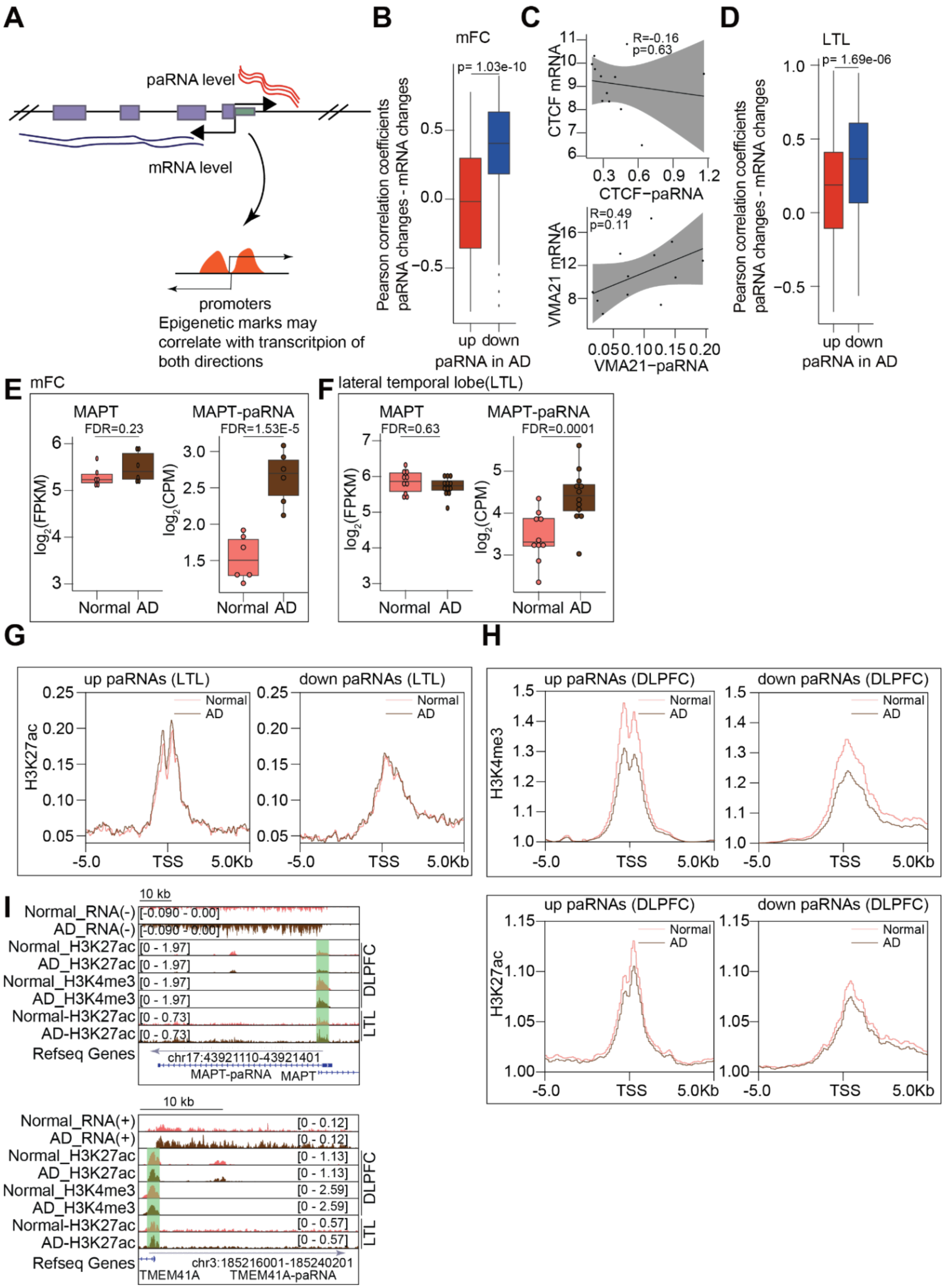
A
paRNA level
mRNA level
promoters
Epigenetic marks may correlate with transcritpion of both directions
B
mFC
p= 1.03e-10
Pearson correlation coefficients paRNA changes - mRNA changes
up down
paRNA in AD
C
R=-0.16
p=0.63
CTCF mRNA
CTCF−paRNA
R=0.49
p=0.11
VMA21 mRNA
VMA21−paRNA
D
LTL
p= 1.69e-06
Pearson correlation coefficients paRNA changes - mRNA changes
up down
paRNA in AD
E mFC
MAPT
FDR=0.23
MAPT-paRNA
FDR=1.53E-5
log2(FPKM)
log2(CPM)
Normal AD
F lateral temporal lobe(LTL)
MAPT
FDR=0.63
MAPT-paRNA
FDR=0.0001
G
up paRNAs (LTL)
down paRNAs (LTL)
H3K27ac
Normal
AD
TSS
H
up paRNAs (DLPFC)
down paRNAs (DLPFC)
H3K4me3
H3K27ac
I
10 kb
Normal_RNA(-)
AD_RNA(-)
Normal_H3K27ac
AD_H3K27ac
Normal_H3K4me3
AD_H3K4me3
Normal-H3K27ac
AD-H3K27ac
Refseq Genes
DLPFC
LTL
chr17:43921110-43921401
MAPT-paRNA MAPT
Normal_RNA(+)
AD_RNA(+)
chr3:185216001-185240201
TMEM41A TMEM41A-paRNA

**Supplementary Figure 6. The correlation of paRNA/mRNA changes in AD and epigenetic state changes at paRNAs' start sites. A**. A cartoon illustrating the possible impact of epigenetic state (i.e., histone modifications) around the shared promoters can affect the expression levels of paRNAs and mRNAs. **B.** A boxplot showing the distribution of Pearson correlation coefficients (PCC) between each AD-dysregulated paRNA and their promoter-sharing mRNA. Red and blue boxplots represent the PCC distribution for AD-upregulated and downregulated paRNAs, respectively. P values were calculated by a two-tailed non-parametric Wilcoxon–Mann–Whitney test. **C.** Two examples of PCCs between an AD-upregulated paRNA and its nearest mRNA, and between an AD-downregulated paRNA and its nearest mRNA. Each PCC is generated from 12 expression values mixing the Normal and AD mFC RNA-seq data. R values were PCC, and p values were calculated by *cor.test*. **D**. Similar to panel B, boxplots showing the PCC distribution for paRNA/mRNA pairs generated by RNA-seq data in LTL from Nativio et al.[36]. Red and blue boxplots represent AD-upregulated and downregulated paRNAs, respectively. **E-F**. Boxplots showing the expression of the *MAPT* gene and *MAPT-paRNA* between Normal and AD brains, respectively. FDR values were calculated by DESeq2. **G-H**. Metaplots showing the enrichment of H3K27ac and H3K4me3 ChIP-seq signals around the TSSs of upregulated and downregulated paRNAs between Normal and AD brains, respectively. The values in the y-axis in boxplots are m6A ratios (IP/input). **I**. Genomic tracks for H3K27ac and H3K4me3 ChIP-seq signals around the TSSs of two example paRNA in Normal and AD brains. In panels **B, D,E, F**, boxplots indicate the interquartile range with the central line representing the median, and the vertical lines extending to the extreme values in the group.

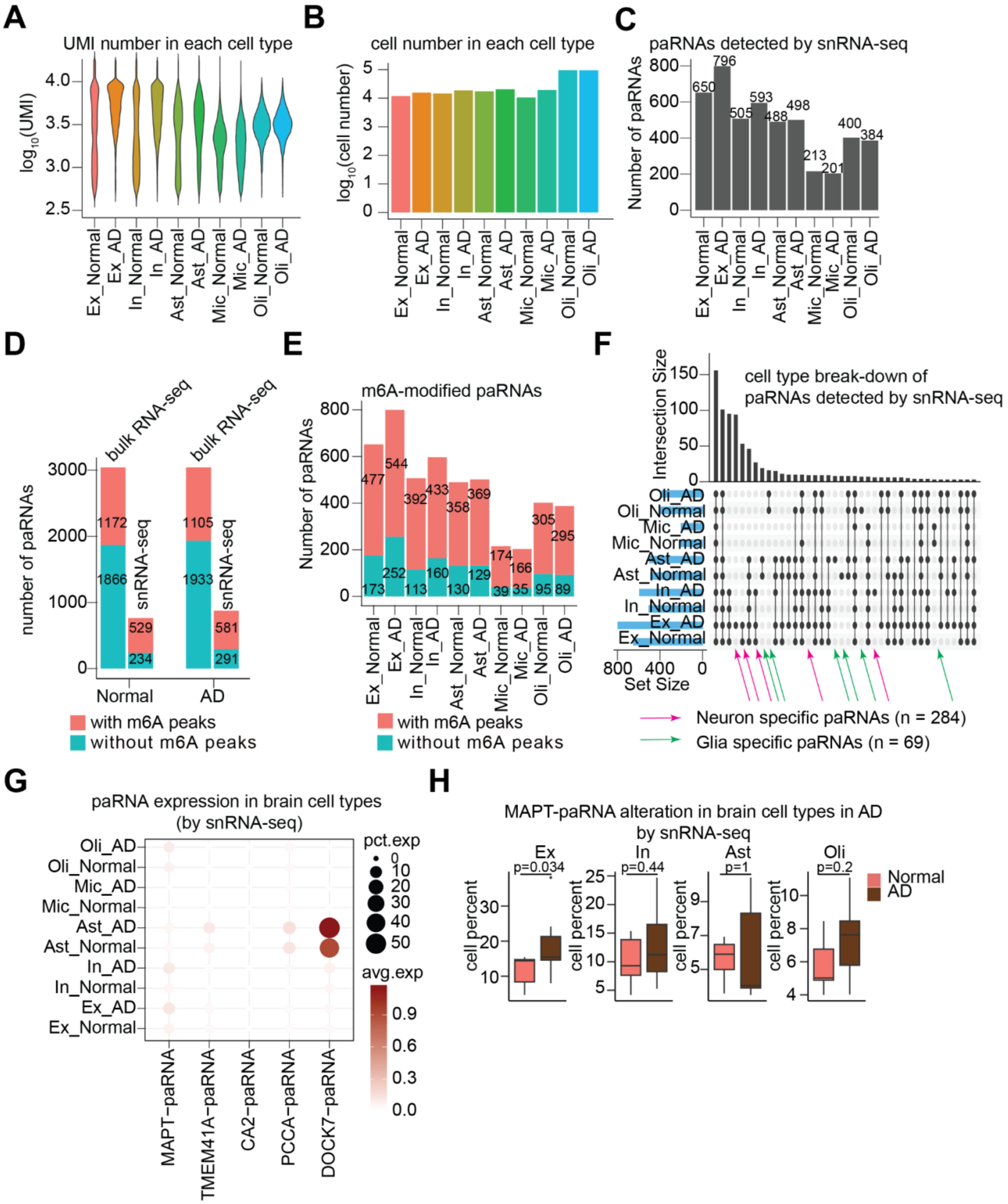

**Supplementary Figure 7. The expression of paRNA at single-cell level in snRNA-seq data. A**. A violin plot showing the numbers of RNA UMIs (unique molecular index) assigned to each cell type in brain samples from snRNA-seq data[52]. **B**. A barplot showing numbers of cells in brain cell types detected by snRNA-seq data[52]. Ex: Excitatory neurons. In: Inhibitory neurons. Ast: Astrocyte. Mic: Microglia. Oli: Oligodendrocyte.**C**. A barplot showing the number of paRNAs detected by single-nuclei RNA-seq data. **D**. A barplot showing the number of paRNAs with m6A peaks or not detected by our bulk RNA-seq and snRNA-seq data[52]. **E**. A barplot showing the number of paRNAs with m6A peaks or not detected by snRNA-seq data. **F**. An UpSet plot showing common and unique paRNAs in different cell types detected by snRNA-seq data from Normal and AD brains. **G**. A dotplot showing the cell-type expression of the top five AD-upregulated paRNAs revealed by snRNA-seq data[52]. Ex: excitatory neurons; In: inhibitory neurons; Ast: astrocytes; Mic: microglia; Oli: oligodendrocytes. **H**. Boxplots showing the percentages of cells of several cell types that express *MAPT-paRNA* by snRNA-seq data[52]. Boxplots indicate the interquartile range with the central line representing the median, and the vertical lines extending to the extreme values in the group. P values were calculated by two-tailed non-parametric Wilcoxon–Mann–Whitney tests.

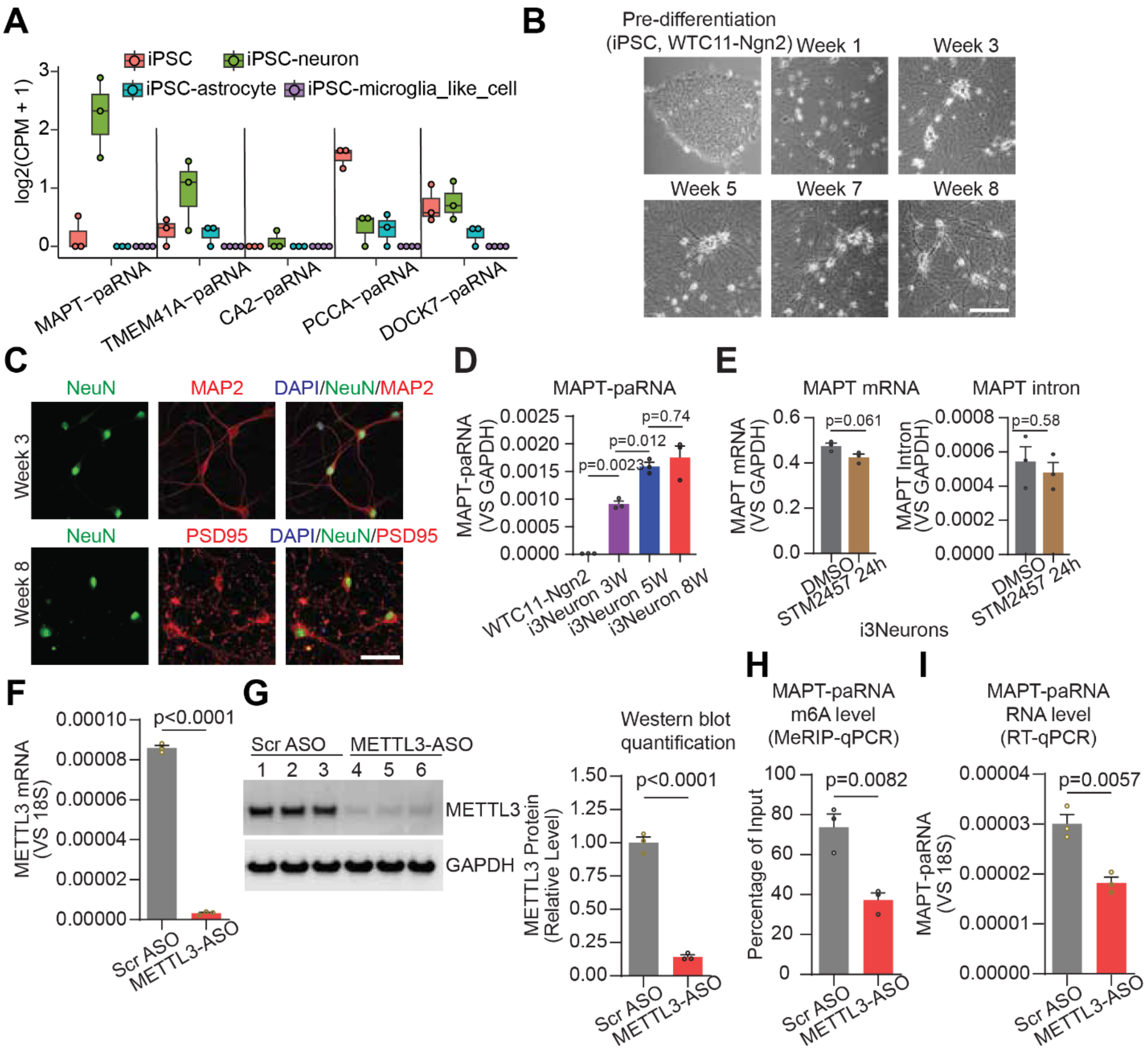


**Supplementary Figure 8. paRNA expression, i3Neuron features and effects of METTL3 inhibition on *MAPT-paRNA* expression A**. Boxplots showing the expression levels of top five AD-upregulated paRNAs revealed by RNA-seq data of iPSC and iPSC-derived neurons, astrocytes and microglia-like cells[53]. Boxplots indicate the interquartile range with the central line representing the median, and the vertical lines extending to the extreme values in the group. **B**. Representative phase-contrast images showing morphological changes of iPSC-derived i3Neurons at different stages of differentiation (from WTC11-Ngn2 induced model). Scale bar, 100µm. **C**. Representative images of immunofluorescent staining of week-3 or week-8 i3Neurons for MAP2 (a neuronal dendrite marker), NeuN (a pan-neuron nuclear marker), and PSD95 (a postsynaptic marker). Scale bar, 100µm. **D**. A barplot showing the expression levels of *MAPT-paRNA* at different stages of i3Neuron differentiation. **E**. Barplots showing the expression levels of *MAPT* gene after *MAPT-paRNA* KD. Data is generated by RT-qPCR using primers targeting either mRNA (across exons) or introns. **F**. A Barplot showing the expression levels of *METTL3* gene mRNA after *METTL3* KD. Data is generated by RT-qPCR using primers targeting *METTL3* mRNA. **G**. (Left) The protein level of *METTL3* in i3Neurons with/without *METTL3* knockdown revealed by western-blot analysis. (Right) Barplot shows the relative RNA expression of *METTL3* quantified by RT-qPCR after a 5-day knockdown of *METTL3*. **H-I**. MeRIP-qPCR and RT-qPCR data showing m6A methylation and *MAPT-paRNA* expression with and without *METTL3* knockdown, respectively. P values were calculated by a two-tailed Student's t-test.

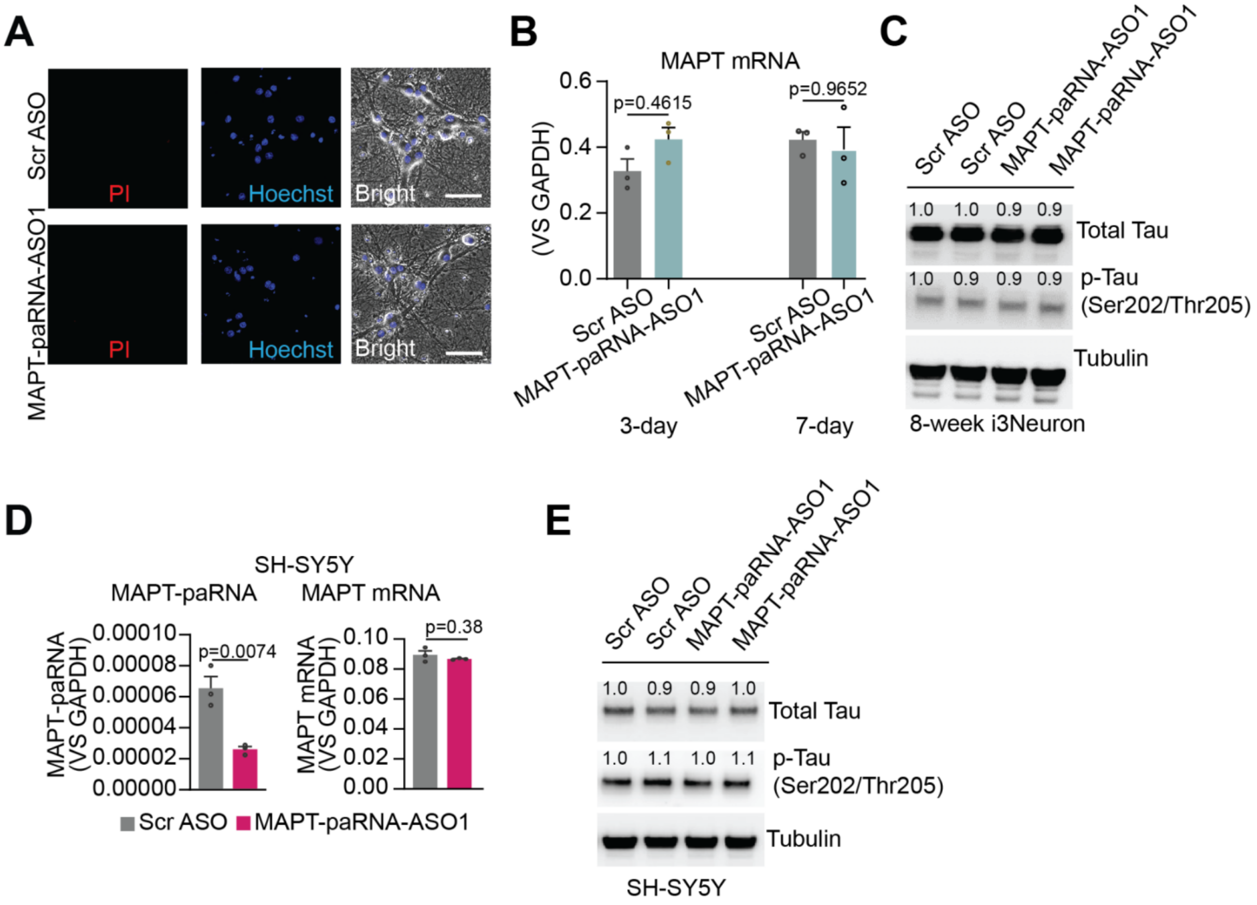


**Supplementary Figure 9. *MAPT-paRNA* does not regulate *MAPT* expression at mRNA or protein levels. A**. Representative images demonstrating that *MAPT-paRNA* knockdown by ASO1 does not affect neuronal viability or morphology. Scale bar = 50 µm. **B**. RT-qPCR analysis of *MAPT* mRNA expression levels after 3 or 7 days of treatment with scrambled ASO or *MAPT-paRNA* ASO1 in i3Neurons. **C**. Western blot analysis (duplicates) showing no significant change in total Tau or phosphorylated Tau (p-Tau) protein levels following 3 days of *MAPT-paRNA* ASO1 treatment in i3Neurons. **D**. RT-qPCR analysis showing no significant change in *MAPT* mRNA expression after *MAPT-paRNA* knockdown by ASO1 in SH-SY5Y cells (a neuroblastoma cell line). **E**. Western blot analysis (duplicates) showing total Tau or phosphorylated Tau (p-Tau) protein levels following 3 days of *MAPT-paRNA* ASO1 treatment in SH-SY5Y cells. The numbers in the panel of C and E represent relative expression levels. P values were calculated by a two-tailed Student's t-test.

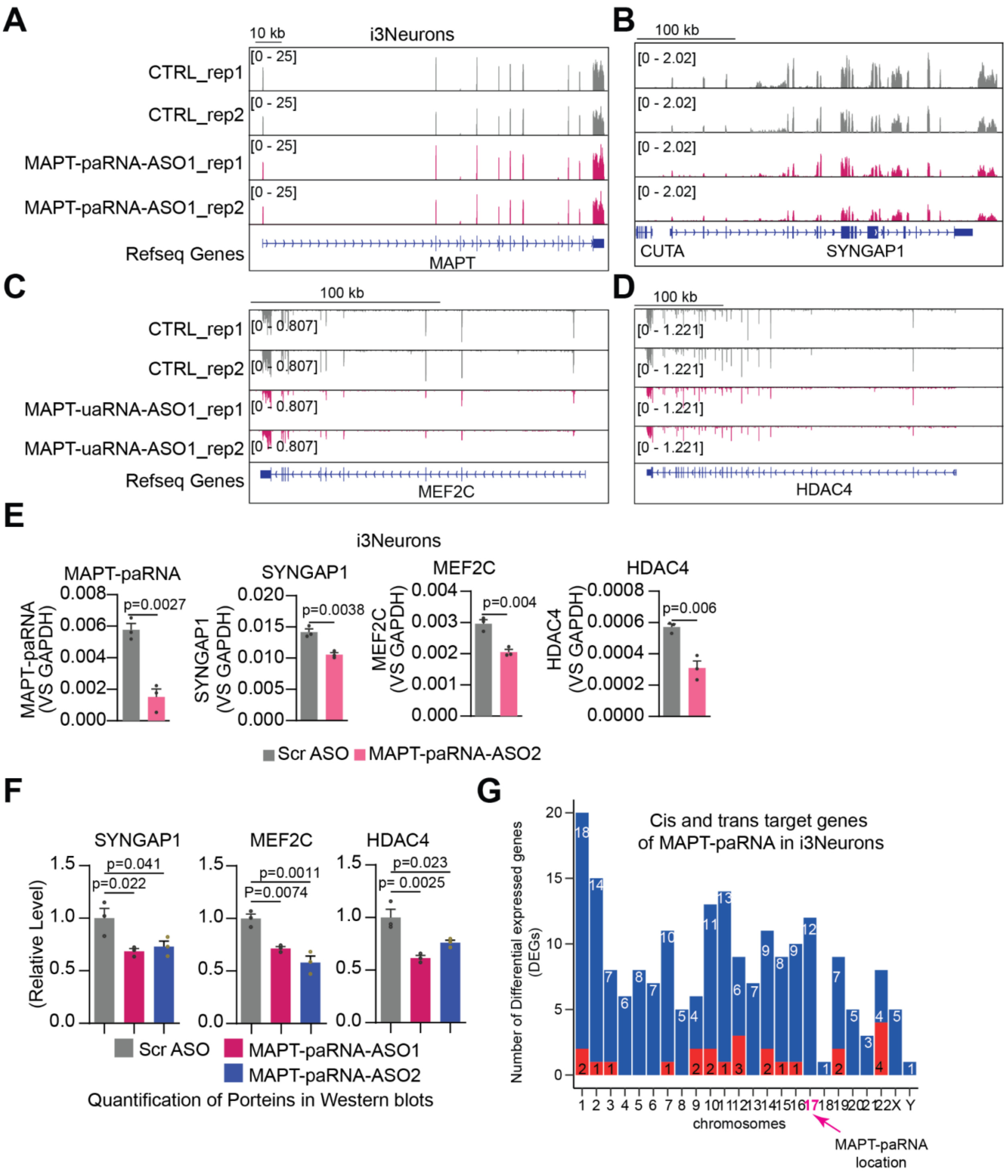


**Supplementary Figure 10.** ***MAPT-paRNA* synaptic gene expression. A-D**. Genomic tracks for *MAPT, SYNGAP1, MEF2C* and *HDAC4* expression in 8 weeks old i3Neurons before and after *MAPT-paRNA* knockdown. **E**. Barplots showing the relative RNA expression of *MAPT-paRNA, MEF2C, SYNGAP1*, and *HDAC4* quantified by RT–qPCR after *MAPT-paRNA* knockdown by a second ASO. **F**. Barplots showing quantification of the WB bands intensity (**Fig. 4I**). **G**. A barplot showing the chromosomal locations of differentially expressed genes after *MAPT-paRNA* knockdown in i3Neurons. Red and blue bars represent upregulated and downregulated genes, respectively. A pink arrow points to the chromosome 17, where *MAPT-paRNA* is transcribed. P values were calculated by a two-tailed Student's t-test.

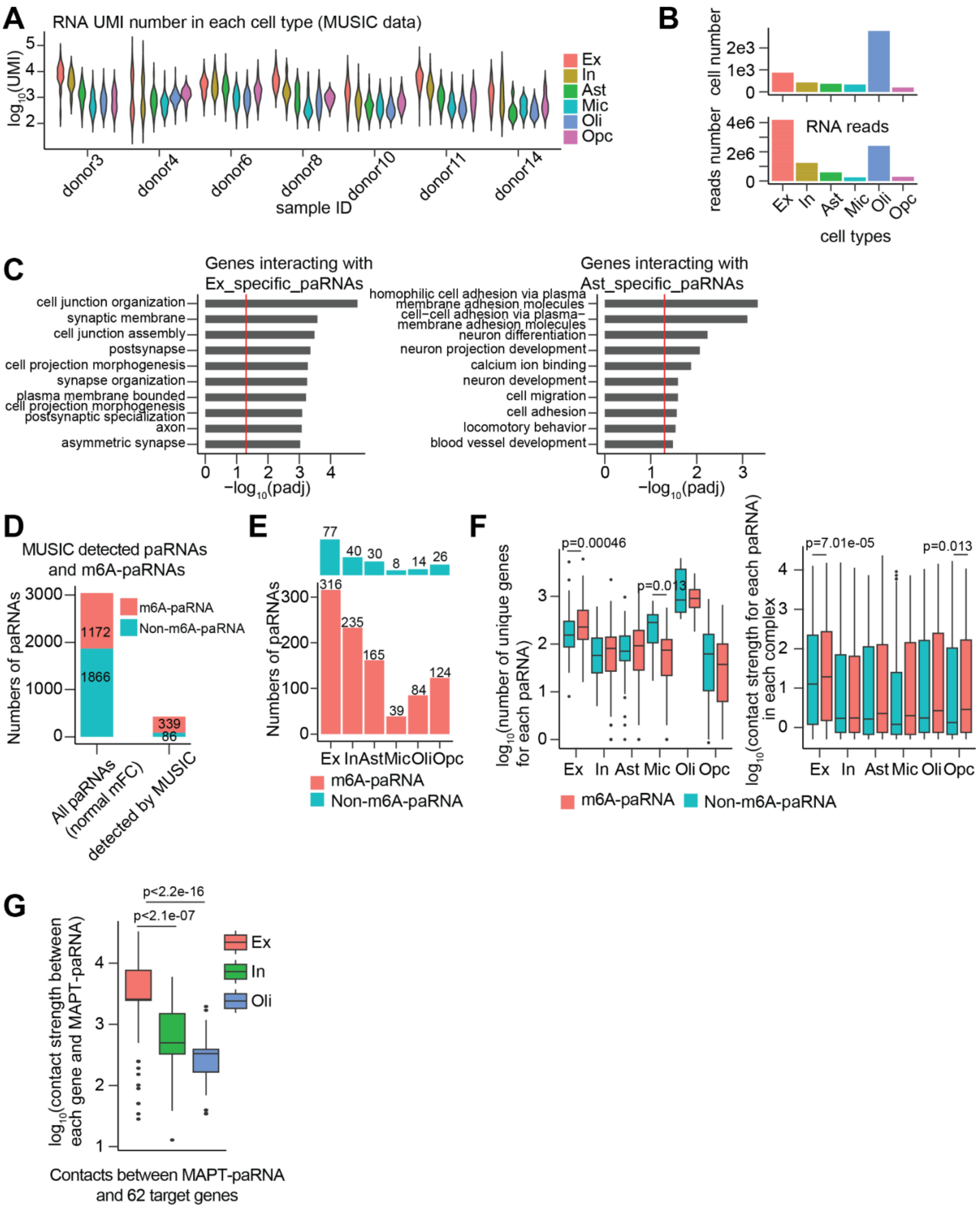


**Supplementary Figure 11. Single cell RNA-DNA interactome from MUSIC to study paRNA-DNA interactions. A**. A violin plot showing the numbers of RNA UMIs (unique molecular index) assigned to each cell type in brain samples from 7 individual donors of the MUSIC dataset[59]. **B**. Barplots showing numbers of cells and RNA sequencing reads in brain cell types detected by MUSIC data[59]. **C**. GO analyses for the target genes of two groups of paRNAs detected to be excitatory neuron specific and astrocyte specific (as in **Fig. 5B**). **D**. A barplot showing detected numbers of m6A-paRNAs and non-m6A-paRNAs by MUSIC data in any cell types. **E**. A barplot showing detected m6A-paRNAs and non-m6A-paRNAs in different cell types by MUSIC data. **F**. (Left) a boxplot showing unique target genes bearing RNA-DNA contacts with m6A-paRNAs or non-m6A-paRNAs detected by MUSIC data in each cell type, and (Right) a boxplot showing the contact strength between target genes and m6A-paRNAs or non-m6A-paRNAs detected by MUSIC data in each cell type. **G**. A boxplot showing the contact strength between *MAPT-paRNA* and its 62 target genes in three cell types that display *MAPT-paRNA* expression. P values were calculated by a two-tailed non-parametric Wilcoxon–Mann–Whitney test. Boxplots indicate the interquartile range with the central line representing the median, and the vertical lines extending to the extreme values in the group.

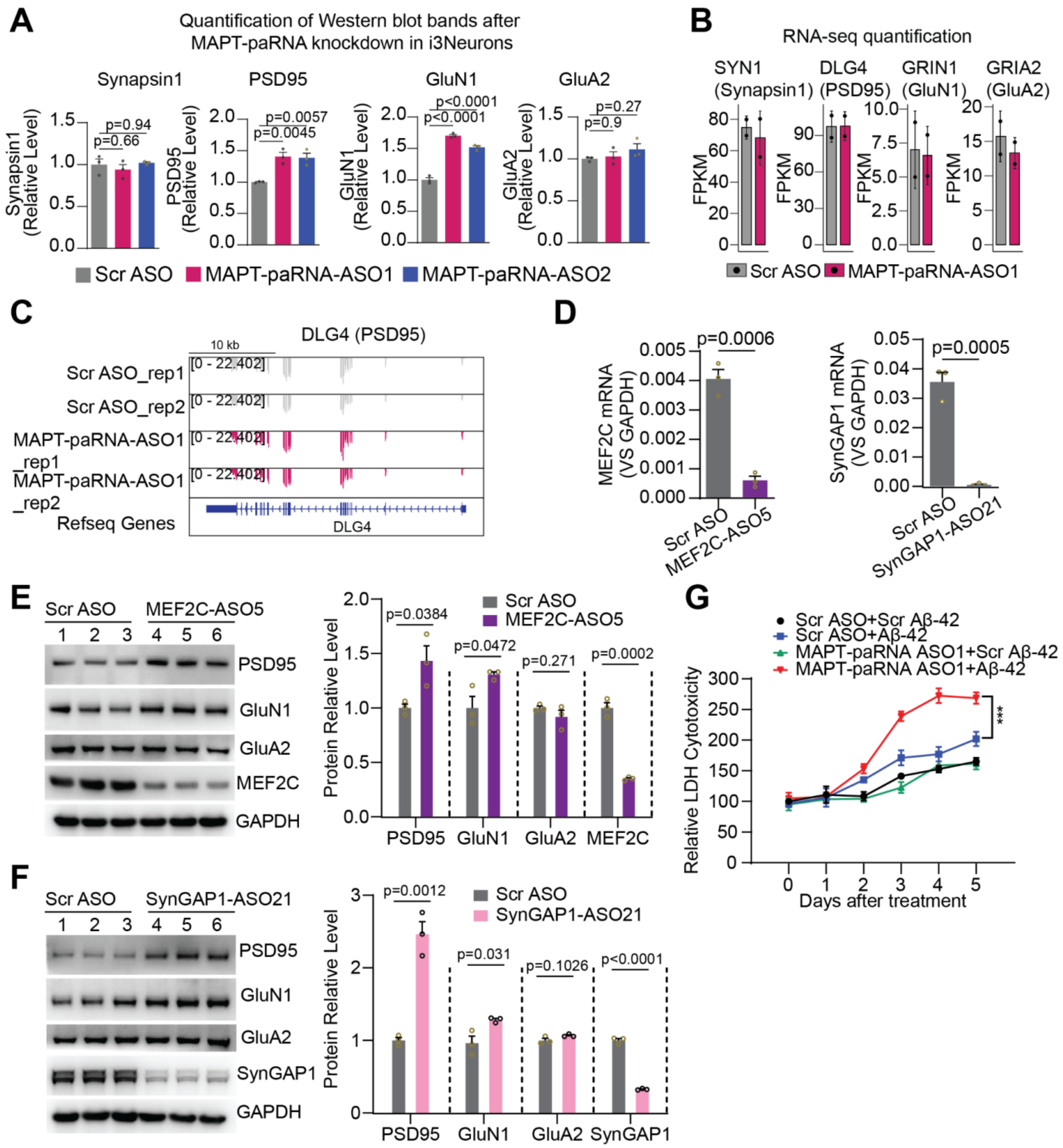


**Supplementary Figure 12. Additional data on *MAPT-paRNA* and its target genes MEF2C and SynGAP1 in regulation of postsynaptic proteins, glutamate receptors and neuronal survival. A**. Quantification of Western blot band intensities for Synapsin1, PSD95, GluN1, and GluA2 following MAPT-paRNA knockdown by ASO1 or ASO2 in i3Neurons. **B**. RNA-seq quantification of *SYN1*, *DLG4*, *GRIN1*, and *GRIA2* expression levels after MAPT-paRNA knockdown by ASO1 in i3Neurons. **C**. Genomic tracks displaying RNA-seq analysis of DLG4 expression in i3Neurons following 3 days of treatment with scrambled ASO (CTRL) or *MAPT-paRNA* ASO1. **D**. RT-qPCR showing the reduction in *MEF2C* or *SynGAP1* mRNA expression after knockdown by ASOs in i3Neurons. **E-F**. Western blot and corresponding quantification of postsynaptic marker PSD95 and glutamate receptors (GluN1 and GluA2) in i3Neurons after *MEF2C* (**E**) or *SynGAP1* (**F**) knockdown. **G**. *MAPT-paRNA* knockdown exacerbates beta-amyloid (Aβ-42)-induced cytotoxicity. Cultured i3Neurons were treated with either Scr ASO or MAPT-paRNA-ASO1 in the presence of scrambled beta-amyloid (Scr Aβ-42) or beta-amyloid (Aβ-42) for six consecutive days. Cell cytotoxicity was evaluated daily by measuring lactate dehydrogenase (LDH) levels in the culture media. P values were calculated by a two-tailed Student's t-test for **A**, **D**, **E**, and **F**; Two-way ANOVA with Tukey's multiple comparisons test for **G** (***, $p<0.0001$).
